# R-loop landscapes in the developing human brain are linked to neural differentiation and cell-type specific transcription

**DOI:** 10.1101/2023.07.18.549494

**Authors:** Elizabeth A. LaMarca, Atsushi Saito, Amara Plaza-Jennings, Sergio Espeso-Gil, Allyse Hellmich, Michael B. Fernando, Behnam Javidfar, Will Liao, Molly Estill, Kayla Townsley, Anna Florio, James E. Ethridge, Catherine Do, Benjamin Tycko, Li Shen, Atsushi Kamiya, Nadejda M. Tsankova, Kristen J. Brennand, Schahram Akbarian

## Abstract

Here, we construct genome-scale maps for R-loops, three-stranded nucleic acid structures comprised of a DNA/RNA hybrid and a displaced single strand of DNA, in the proliferative and differentiated zones of the human prenatal brain. We show that R-loops are abundant in the progenitor-rich germinal matrix, with preferential formation at promoters slated for upregulated expression at later stages of differentiation, including numerous neurodevelopmental risk genes. RNase H1-mediated contraction of the genomic R-loop space in neural progenitors shifted differentiation toward the neuronal lineage and was associated with transcriptomic alterations and defective functional and structural neuronal connectivity *in vivo* and *in vitro*. Therefore, R-loops are important for fine-tuning differentiation-sensitive gene expression programs of neural progenitor cells.

## INTRODUCTION

Transcriptional regulation is developmental stage- and tissue-specific, with fine-tuning of sequentially organized gene expression programs during the course of neuronal and glial differentiation critical for normal brain function and behavior[1–3]. However, some mechanisms that have a fundamental impact on transcriptional regulation still lack genome-scale exploration in the developing brain. This includes R-loops, three-stranded structures consisting of a DNA/RNA hybrid and a displaced single strand of DNA [4, 5]. R-loops can interfere with DNA methyltransferase-mediated 5mC deposition over CpG-island promoters [4], regulate transcription factor binding at gene regulatory elements [6–8], and can aid in efficient transcription termination when forming over distal genic sequences [9]. On the other hand, emergence of unscheduled R-loops within the genome can trigger DNA damage, transcription-associated recombination [10, 11], and can foster replication-collision events in dividing cells [12]. Therefore, formation and resolution of R-loops requires a delicate balance in order to effect transcriptional regulation while maintaining genome stability.

As much as 2-12% of the genome can assemble as R-loops, most frequently at RNA polymerase II-transcribed sequences, promoters, gene bodies, and subsets of repeat-associated intergenic sequences associated with chromatin modifications that facilitate gene expression [13]. To date, no genome-scale R-loop mappings exist for the nervous system, including human. Little is known about the potential roles of R-loops for developmentally regulated gene expression programs in the brain, although an increasing list of monogenic brain disorders has been linked to DNA mutations and structural variants in enzymes that resolve R-loop structures, including DNA topoisomerases [14] and RNA nucleases and helicases including RNase H [15] and senataxin [16, 17]. In addition, R-loops have been invoked in the molecular pathophysiology of triplet and other repeat expansion-based neurological diseases including Fragile-X syndrome, Friedreich’s ataxia, and C9orf72-mediated amyotrophic lateral sclerosis [18–20].

Here, we generated high-resolution maps of R-loop distribution during human neurodevelopment using prenatal postmortem and *in vitro* human induced pluripotent stem cell (hiPSC)-derived models, focusing on developmental transitions from neural progenitor to a more fully differentiated stage. We report that R-loops cover approximately 2% of the genome in neural progenitor cells (NPCs) and are highly dynamic across subsequent neuronal differentiation. Experimentallyinduced contraction of the R-loop landscape severely disrupts the temporal tuning of gene expression on a genome-wide scale, giving rise to premature expression of many neuron-specific transcripts at sites of promoter-bound R-loops, resulting in lasting defects in functional and structural neuronal connectivity.

## RESULTS

### Genome-scale R-loop mapping in proliferative and differentiated zones of the prenatal human cerebral cortex

Genome-scale R-loop studies most commonly employ the monoclonal anti-DNA/RNA hybrid antibody, S9.6. We therefore first conducted a series of quality control assays. We confirmed the specificity of S9.6 for DNA/RNA hybrids relative to double-stranded DNA (dsDNA) and dsRNA (Supplementary Figure 1A). We then replicated in our NPCs two findings previously reported in peripheral cells, including punctate intranuclear staining patterns with S9.6 immunocytochemistry (Supplementary Figure 1B) [21], and an approximately 50% reduction of R-loop occupancies genome-wide when genomic neural progenitor cell (NPC) DNA was treated with an RNase H1/A cocktail prior to S9.6 DNA/RNA Immuno-Precipitation followed by next generation sequencing (DRIP-seq) [22, 23] (Supplementary Figure 1C,D). Further, NPC RNA alone was taken through the DRIP-seq protocol as opposed to genomic DNA, which produced few DRIP-seq peaks (n = 53), thereby establishing that dsRNA or single-stranded (ss)RNA makes a negligible contribution to the peaks called in a DRIP-seq assay (Supplementary Figure 1D).

We then assayed R-loops from two anatomical regions of the human prenatal post-mortem brain during the second trimester (17-21 gestational week), including the germinal matrix (GM, comprised primarily of mitotically active NPCs, n = 3), and cortical plate (CP, comprised primarily of post-mitotic neurons, n = 3) (Figure 1A, Supplementary Table 1). DRIP-Seq libraries for prenatal GM and prenatal CP did not significantly differ in number of peaks (GM average ± SEM: 52,686 ± 11,840; CP: 70,199 ± 8,043) and peak length (GM average ± SEM: 1,062 bp ± 85; CP: 969 ± 32), covering in total an average of 63 Mbp of genome for each sample (One-way ANOVA with Tukey’s Multiple Comparison test; Supplementary Figure 2A- E; Supplementary Table 2). Of note, R-loops showed a distinct landscape among brain regions (Figure 1B), suggesting strong cell type- and/or developmental stage-specific roles of R-loops in the brain. We identified 11,612 DRIP-seq peaks selectively enriched in GM (FDR < 0.01, log_2_ fold change > 2.2), and 7,685 DRIP-seq peaks selectively enriched in the CP (log_2_ fold change < -2.2) (Supplementary Table 3). Differentially regulated R-loops showed robust functional signatures corresponding to the differentiation status of each of the two brain regions: top gene ontology (GO) terms for genes at sites of GM-specific R-loops were linked to embryo and cell development, cell projection and cytoskeletal organization, and neurogenesis. In contrast, genes at sites of CP-specific R-loops showed significant enrichment for cell adhesion, synapse assembly, transmission, and signaling, forebrain neuron development and other neuronal functions (Figure 1C; Supplementary Tables 4-5).

**Figure 1.**
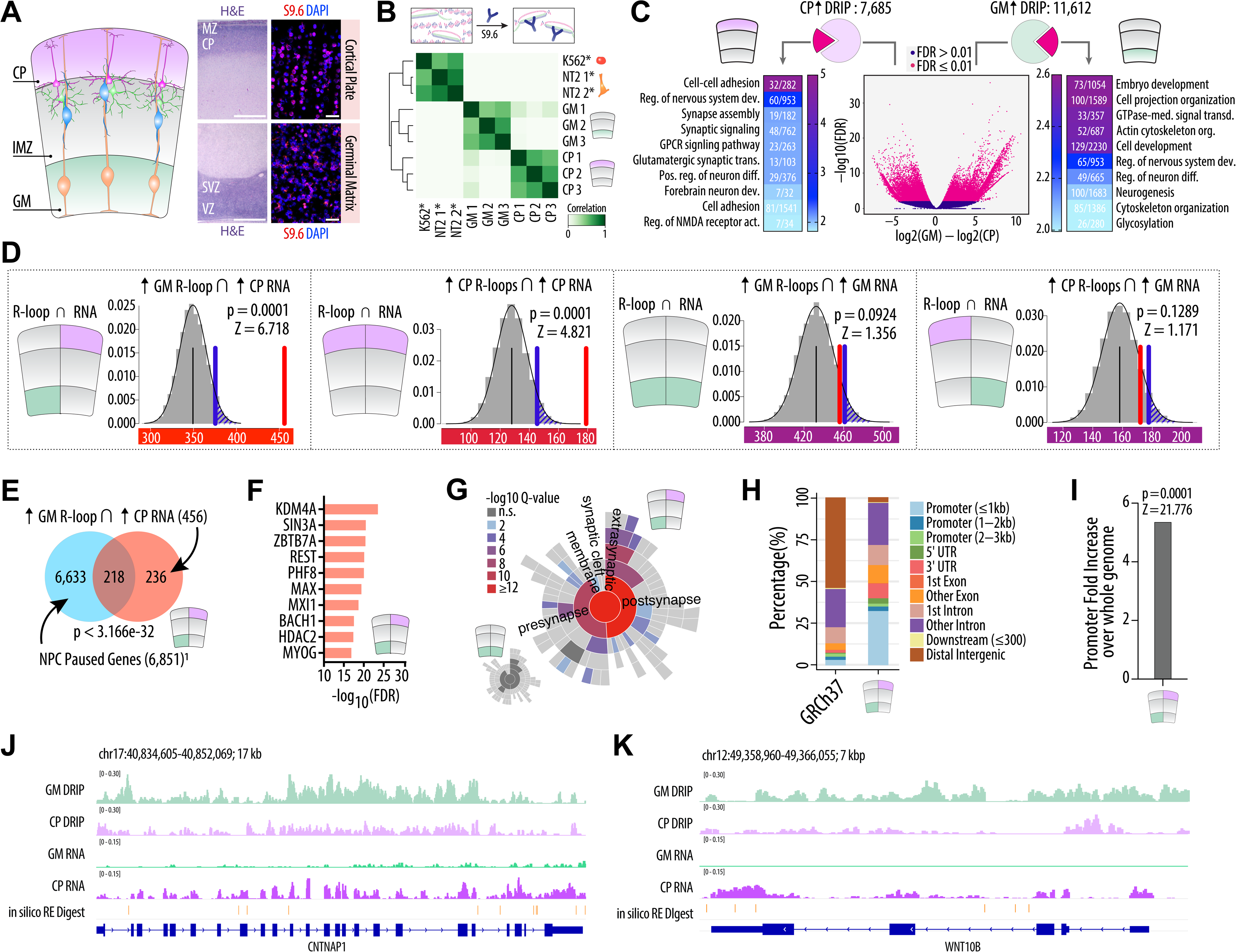
DRIP-seq in prenatal brain and neural cells in vitro. **A)** Schematic of the second trimester prenatal brain, containing the cortical plate (CP) and the germinal matrix (GM). Left panel: H&E stain of the CP, located under the marginal zone (MZ), and the GM, containing the subventricular and ventricular zones (SVZ, VZ); scale bar: 1 mm. Right panel: Immunofluorescent staining with the R-loop-specific antibody S9.6 (red) and the nuclear counterstain DAPI (blue) in each respective brain region; scale bar: 20 mm. **B)** Heatmap showing DRIP-seq peaks across prenatal CP & GM and non-neuronal cells NTera2 and K562 (* = data from GSE70189 [5]), indicating R-loops cluster by cell type and brain region. **C)** Center: Volcano plot displaying the 37,698 differential DRIP-seq peaks between prenatal GM (n = 3) and CP (n = 3) (FDR < 0.01, pink dots). The program DiffBind [99] was used to find differential R-loop peaks between GM and CP. Left panel: top GO terms for the 7,685 CP-enriched R-loops forming over 5,182 unique genes (log_2_ fold change < -2.2). Right panel: top GO terms for the 11,612 GM-enriched R-loops forming over 9,106 unique genes (log_2_ fold change > 2.2). Numbers within GO heatmaps: Brain region-enriched genes in GO category / total genes in GO category; scale bar: -log_10_ FDR. **D)** Top left: The number of overlapping genes between GM-enriched R-loops and CP-enriched DEGs (BrainSpan RNA microarray data [24]) (red bar) (overlapping n = 456) was tested for significance by performing 10,000 random permutations of the overlap of GM-enriched R-loops with a “background universe” of any expressed gene in the prenatal cortex (gray bars comprising normal distribution), blue bar = significance threshold (α = 0.05). Remaining graphs similarly summarize permutation results for CP-enriched R-loops ∩ CP-enriched DEGs (second from left; overlapping n = 179), GM-enriched R-loops ∩ GM-enriched DEGs (second from right; overlapping n = 456), and CP-enriched R-loops ∩ GM-enriched DEGs (right; overlapping n = 172). Positive Z-scores indicate overlap of the target gene sets are higher than expected by chance. **E)** A significant proportion of genes in the set GM-enriched R-loops ∩ CP-enriched DEGs (overlap n = 218, p < 3.166e-32) have previously been defined as NPC-specific poised genes (see main text for more detail). **F)** Transcription factors (ENCODE) [100] that target genes in the sets GM-enriched R-loops ∩ CP-enriched DEGs. **G)** A significant proportion of genes in the set GM-enriched R-loops ∩ CP-enriched DEGs belong to synaptic gene ontology within the SynGO [31] package (82 of 456). Small inset image (bottom left) depicts SynGO enrichment in the set GM-enriched R-loops ∩ GM-enriched DEGs (22 of 456, not significant). **H-I)** R-loops within the set GM-enriched R-loops ∩ CP-enriched DEGs are significantly enriched in promoters relative to genomic GRCh37 background (5.39-fold enriched). **J-K)** Browser tracks over two putatively R-loop primed genes, *CNTNAP1* and *WNT10B* (top to bottom) DRIP-seq R-loop signal in the prenatal GM and CP, RNA-seq GM and CP. Orange track, *in silico* HindIII/EcoRI/XbaI/ SSPI restriction digest (restriction enzyme cocktail used in DRIP-seq protocol), for resolution of DRIP-seq peaks.

### R-loops are enriched in the germinal matrix over poised gene promoters

R-loops are broadly associated with gene expression, occurring co-transcriptionally with site-specificity and highly dynamic turnover [5]. We therefore asked whether R-loop regulation in the prenatal human brain is linked to region-specific differences in gene expression, between GM, as a proliferative, and CP, as a differentiated, neuron-specific layer of the developing cortex. To this end, we compared our R-loop maps in the prenatal GM and CP with the patterns of GM- and CP-specific gene expression, using the BrainSpan resource [24] with focus on 16-21 GW fetal brains matching the age range of our brains used for DRIP-seq.

Consistent with the general association of R-loops with the transcriptional process, there was a significant association with the set of CP-specific differentially expressed genes (DEG) and CP-specific DRIP-seq datasets (10,000 random permutations, Z = 4.821, p = 0.0001, Figure 1D). These findings were tissue specific because the genomic R-loop space of neither NTera2 embryonic carcinoma nor K562 lymphoblasts ([5], GSE70189) revealed any significant association CP or GM-specific genes (Supplementary Figure 3A, B). Unexpectedly, however, GM-enriched R-loops robustly and significantly overlapped with genes transcriptionally upregulated in the CP, but not genes upregulated in the GM (Figure 1D: GM-enriched R-loops ∩ CP-enriched DEGs, overlap n = 456, Z = 6.718, p = 0.0001; GM-enriched R-loops ∩ GM- enriched DEGs, overlap n = 456, Z = 1.356, p = 0.0924; Supplementary Figure 3A; Supplementary Table 6). These results suggest that a subset of GM-specific R-loops are positioned at genes programmed for increased expression during subsequent neuronal differentiation. To further test this hypothesis, we examined a group of 6,633 genes previously identified as primed (programmed for transcriptional up-regulation during later stages of differentiation) in neural progenitor cells (the defining cell type in the GM) from embryonic E15.5 mouse brain [25]. Indeed, the group of N = 456 GM-specific R-loops at sites of differentiation-sensitive genes (GM-enriched R-loops ∩ CP-enriched DEGs) from our human brains showed a significant overlap with genes primed for upregulation in mouse neural progenitor cells (Orthologue-converted R-loop-primed genes ∩ NPC-primed genes, overlap n = 218, p < 3.166e-32 (Figure 1E); hypergeometric test).

Next, we assessed regulatory motifs (ENCODE) for the group of our N = 456 GM-specific R-loops at differentiation-sensitive genes (GM-enriched R-loops ∩ CP-enriched DEGs) and found that the top-ranking transcription factor targets of these R-loop-primed gene sequences are involved in multiple types of repressor functions, including the histone lysine demethylase KDM4A [26], the ZBTB7A zinc finger protein [27], the SUZ12 polycomb repressive complex subunit [28], and the SIN3A [29] and REST [30] transcriptional repressors (Figure 1F). Furthermore, as expected for genes primed for transcriptional upregulation in differentiated CP neurons, this group of R-loop-primed GM genes showed significant enrichment in neuronal synapse ontology (GM-enriched R-loops ∩ CP-enriched DEGs: 88 genes of 454 in the synaptic gene set, -log_10_ Q-value ≥ 12; SynGO, Fisher’s exact test, all brain-expressed genes from GTEx v7 [31] as background set) (Figure 1G, Supplementary Table 7). Notably, R-loops within these putatively primed genes showed a significant, 5.4-fold increase over promoter regions relative to the GRCh37 background (Figures 1H, 1I). Figures 1J-K compare GM- and CP-specific DRIP-seq and RNA-seq profiles over two representative neurodevelopmental loci harboring R-loop primed genes, *CNTNAP2* [32] and *WNT10B* [33, 34]. Taken together, our findings strongly suggest that the genome of immature cells located in the proliferative layers of the prenatal brain harbors R-loops over several hundred genes that are slated for increased expression after neurons differentiate and settle in the CP. We therefore wondered whether this subgroup of primarily promoter-bound R-loops could contribute to transcriptional poising in *cis*, thereby fine-tuning the precise timing of neuronal gene expression during the transition from progenitor status in development.

### Global R-loop decrease in neural precursors promotes neuronal differentiation and upregulated expression of neuronal genes with promoter-bound R-loops

To test our hypothesis that R-loops indeed prime gene expression programs in the developing human brain, we next sought to experimentally contract the genomic R-loop space in our *in vitro* neural precursor cell (NPC) system (Figure 2A, Supplementary Table 8). Of note, the transcriptome of cultured NPCs is closely related to the gene expression program of the GM in the fetal brain [35, 36]. Because RNase H1 specifically destroys the RNA moiety of DNA/RNA hybrids [37] and has been successfully employed in peripheral cell lines to experimentally knock down R-loops [38–42], we designed a doxycycline-inducible, catalytically active RNase H1 lentiviral overexpression system, in addition to a mutant (D145N) RNase H1, which substantially reduces RNase H1 catalytic activity [43]. The mitochondrial localization signal (MLS) was removed from both of these RNase H1 cDNAs, as degradation of mitochondrial R-loops has been shown to be deleterious to cell health [44] (Figure 2B-C, Supplementary Figure 4A). We transduced hiPSC-NPCs with the TRE3G-*RNASEH1*ΔMLS-PGK-BFP-PuroR construct (hereafter referred to as RH1ΔMLS) or the TRE3G-*RNASEH1*ΔMLS^D145N^-PGK-BFP-PuroR construct (hereafter referred to as RH1ΔMLS^D145N^). As an additional negative control, cells were also transduced with the lentiviral backbone containing BFP alone (Figure 2C).

**Figure 2.**
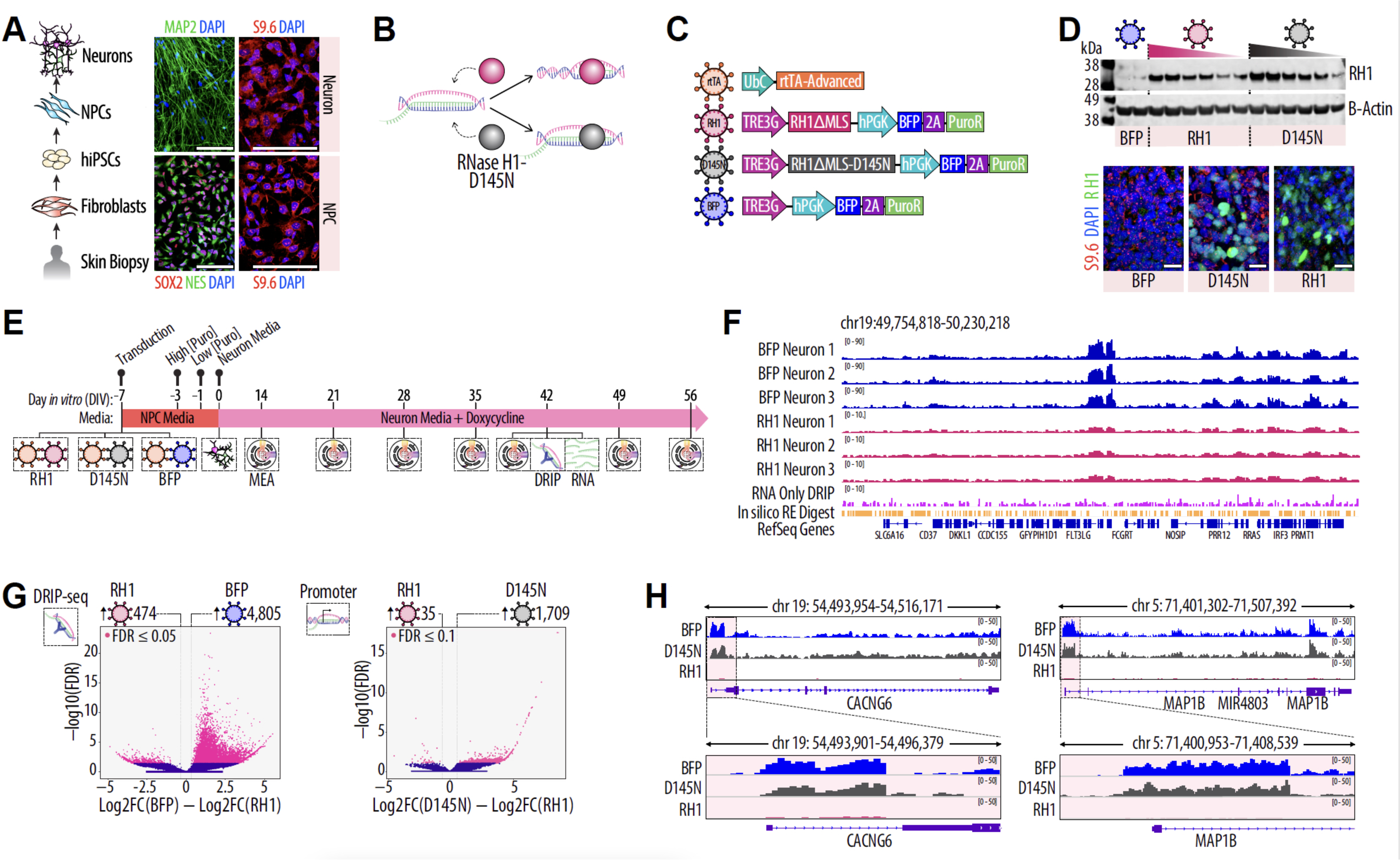
Inducible RNase H1 transgene expression in vitro. **A)** Schematic of hiPSC differentiation from fibroblasts to NPCs and neurons. Left panel: Immunofluorescent staining of the neuronal marker MAP2 (green) in hiPSC-neurons (top) and the NPC markers SOX2 (green) and Nestin (red) in hiPSC-NPCs (bottom); scale bar: 100 mm. Right panel: Immunofluorescent staining of S9.6 (red) and DAPI (blue) in each respective cell type; scale bar: 100 mm. **B)** Schematic of the effect of RNase H1-mediated knockdown of R-loops, or D145N mutant that is binding- competent but catalytically inactive. **C)** Overview of the lentiviral constructs used in this study, including the rtTA transactivator, RH1ΔMLS, RH1ΔMLS^D145N^ catalytically-inactive mutant, and BFP-expressing lentiviral backbone control. **D)** (Top) Representative Western Blot of hiPSC-neurons expressing either BFP, RH1ΔMLS, or RH1ΔMLS^D145N^ lentiviruses. Gradient triangles indicate concentration of virus. RNase H1 protein is 5.1-fold increased in RH1ΔMLS cells relative to BFP controls (normalized to β-actin loading control). (Bottom) Immunocytochemistry for S9.6 (red) and RNase H1 (green) in hiPSC-neurons expressing BFP, RH1ΔMLS^D145N^, or RH1ΔMLS transgenes. RNase H1 protein is robustly increased in the cell nucleus (DAPI, blue) in RH1ΔMLS and RH1ΔMLS^D145N^ cells relative to BFP controls, indicating successful nuclear expression of the transgene-encoded protein; scale bars = 20 mm. **E)** Experimental timeline. NPCs are transduced with RH1ΔMLS or a control (RH1ΔMLS^D145N^ or BFP backbone) lentivirus. After 4 days *in vitro*, puromycin (1:1,000) is added to the culture media to select for transgene-expressing cells. After 6 days in culture, puromycin concentration is dropped to 1:5,000. On day 0 (indicating transition from NPC to neuron differentiation), cells are replated and fed every 2 days with neuron differentiation medium containing 1:1,000 doxycycline to activate the transgene. Every week from week 2, electrophysiological recordings are captured by multielectrode array (MEA). After 6 weeks in culture, cells are harvested for DRIP-seq, bulk- and scRNA-seq. **F)** (top to bottom) DRIP-seq tracks for hiPSC-derived neurons transduced with BFP (blue tracks, BFP Neuron cultures 1-3) or RH1ΔMLS (bright red tracks, cultures 1-3) constructs over a well-defined, conserved R-loop hotspot region [47], demonstrating robustness of our DRIP-seq experimental and computational processing. Note that y-axis scaling 0-90 for BFP Neuron DRIP-seq is wider than RH1 Neuron scale, 0-10. RNA alone processed by the DRIP-seq protocol (RNA-only DRIP, magenta track) shows no discernable peaks, indicating that our S9.6 immunoprecipitation is specific for DNA/RNA hybrids and does not represent single-stranded or double-stranded RNA. Orange track: *in silico* HindIII/EcoRI/XbaI/ SSPI restriction enzyme digest showing expected cutting sites of the restriction enzyme cocktail used for DRIP-seq. **G)** Left: Volcano plot displaying the 6,380 differential promoter-bound R-loop peaks (DRIP-seq) between hiPSC-differentiated cultures expressing RH1ΔMLS (n = 3) or BFP (n = 3) (FDR < 0.05, pink dots) after 6 weeks in culture. Of these, 4,805 DRIP-seq peaks were enriched in BFP controls (log_2_ fold change > 1) and 474 DRIP-seq peaks were enriched RH1ΔMLS cells (log_2_ fold change < -1), indicating successful knockdown of R-loops in the latter. Right: Volcano plot displaying the 1,753 differential promoter-bound R-loop peaks between RH1ΔMLS (n = 3) and RH1ΔMLS^D145N^ (n = 2) (FDR < 0.1, pink dots) cells after 6 weeks in culture. Of these, 1,709 DRIP-seq peaks were enriched in RH1ΔMLS^D145N^ controls (log_2_ fold change > 1) and 35 DRIP-seq peaks were enriched in RH1ΔMLS cells (log_2_ fold change < -1), again indicating successful knockdown of R-loops in the latter. **H)** Representative DRIP-seq tracks over two neuronal genes for BFP-, RH1ΔMLS^D145N^-, and RH1ΔMLS- expressing hiPSC-neurons after 6 weeks in culture. Pink highlighted region and inset displays magnified promoter region.

hiPSC-NPCs transduced with RH1ΔMLS or RH1ΔMLS^D145N^ constructs showed a robust, up to 5.1-fold increase in RNase H1 protein levels by Western blot relative to BFP controls (Figure 2D, top). mRNA levels of the *RNASEH1* gene were also significantly increased in RH1ΔMLS cells relative to BFP controls (∼2-fold, log scale, p < 0.0001 DESeq2 [45]; Supplementary Figure 4B). Immunocytochemistry confirmed robust intranuclear accumulation of RNase H1 protein following transgene overexpression (Figure 2D, bottom), and DRIP-seq confirmed that the catalytically active RH1ΔMLS transgene specifically reduced R-loop DRIP reads in the nuclear, but not mitochondrial, genome (Supplementary Figure 4C-E). The catalytically active RH1 transgene was overexpressed for the full 42 days of differentiation (Figure 2E), with no apparent loss of viability, consistent with earlier findings in embryonic stem cells [46].

Visual inspection of DRIP-seq tracks in our differentiated hiPSC-neurons expressing wildtype levels of *RNASEH1* over “gold-standard,” conserved R-loop hotspot regions (e.g., the chromosome 19 *FLT3LG/FCGRT* locus [5]), showed the expected peak profile based on a study of DRIP-seq best practices [47] (Figure 2F). Conversely, RH1 transgenic hiPSC-neurons overexpressing catalytically active *RNASEH1* showed a multi-fold decrease in R-loop peak profiles (Figure 2F). Additionally, wildtype neuronal RNA alone was taken through the DRIP-seq protocol and showed no discernable peaks. Therefore, our DRIP-seq/S9.6 immunoprecipitation assay is specific for DNA/RNA hybrids, sensitive to *RNASEH1* dosage variation, and does not capture an appreciable amount of single-stranded or double-stranded RNA (Figure 2F).

As expected, RH1ΔMLS cultures (n = 3 independent cultures) each had significantly fewer R-loop peaks than our five control cultures. RH1ΔMLS cells contained (mean ± SEM) 21,528 ± 940 R-loop peaks, spanning 0.95% ± 0.06% (29.9 ± 1.8 Mbp) of the genome, which represented a 34-57% decrease in peaks relative to controls and a 51-56% contraction of the genome-wide R-loop space in comparison to controls (BFP: p = 0.0061, n = 3; RH1ΔMLS^D145N^: p = 0.0008, n = 2; One-way ANOVA with Tukey’s Multiple Comparison test) (Supplementary Figure 4F-I).

Given that in the prenatal GM, R-loops were present at many promoters poised for differentiation-associated transcriptional upregulation (Figure 1H-I), we examined alterations in promoter-bound (± 300 bp from TSS) R-loop occupancies following RNase H1 transgene expression in our *in vitro* system. We observed a more than 10-fold reduction of promoter-bound R-loops in RH1ΔMLS neurons (4,805 BFP vs. 474 RH1ΔMLS, FDR ≤ 0.05, log_2_ fold change < - 1; 1,709 RH1ΔMLS^D145N^ vs. 35 RH1ΔMLS, FDR ≤ 0.1, log_2_ fold change < -1) (Figure 2G; Supplementary Tables 9-10 for promoters and 11-12 for genome-wide R-loop space; Supplementary Figures 4J-K). Figure 2H shows representative examples of DRIP-seq tracks for control and RH1 transgenic cells after six weeks in culture, over two neurodevelopmental risk loci at chromosomes 19 and 5, namely the voltage-gated Ca^2+^ channel, *CACNG6,* and the cytoskeletal protein and regulator of neuronal connectivity, *MAP1B*, respectively.

Next, we wanted to examine the effects of our genome-scale R-loop knockdown during NPC differentiation on cell specifications and patterns of gene expression. To this end, we performed single-cell RNA-sequencing (scRNA-seq) on cultures differentiated from hiPSC-NPCs. Because binding-competent but catalytically inactive mutants of RNase H1 can interfere with R-loop dynamics and downstream transcriptional activity [48, 49], we limited our transcriptomic assay to BFP cultures as the primary control (as opposed to RH1ΔMLS^D145N^). scRNA-seq differential expression analysis (n = 2 replicates each from 2 donors/group: BFP and RH1ΔMLS) revealed significantly higher *RNASEH1* expression levels in RH1ΔMLS cells than BFP controls, as expected (p < 0.0001; log_2_ fold change = 3.86; Supplementary Table 13), and a dramatic effect of *RNASEH1* overexpression on neuronal gene expression profiles. Specifically, gene set enrichment analysis (GSEA) indicated that RNase H1-overexpression led to increased expression of genes associated with neuron differentiation, neuron projection, neuronal development, and synapse gene ontology categories (Figures 3A-B).

**Figure 3.**
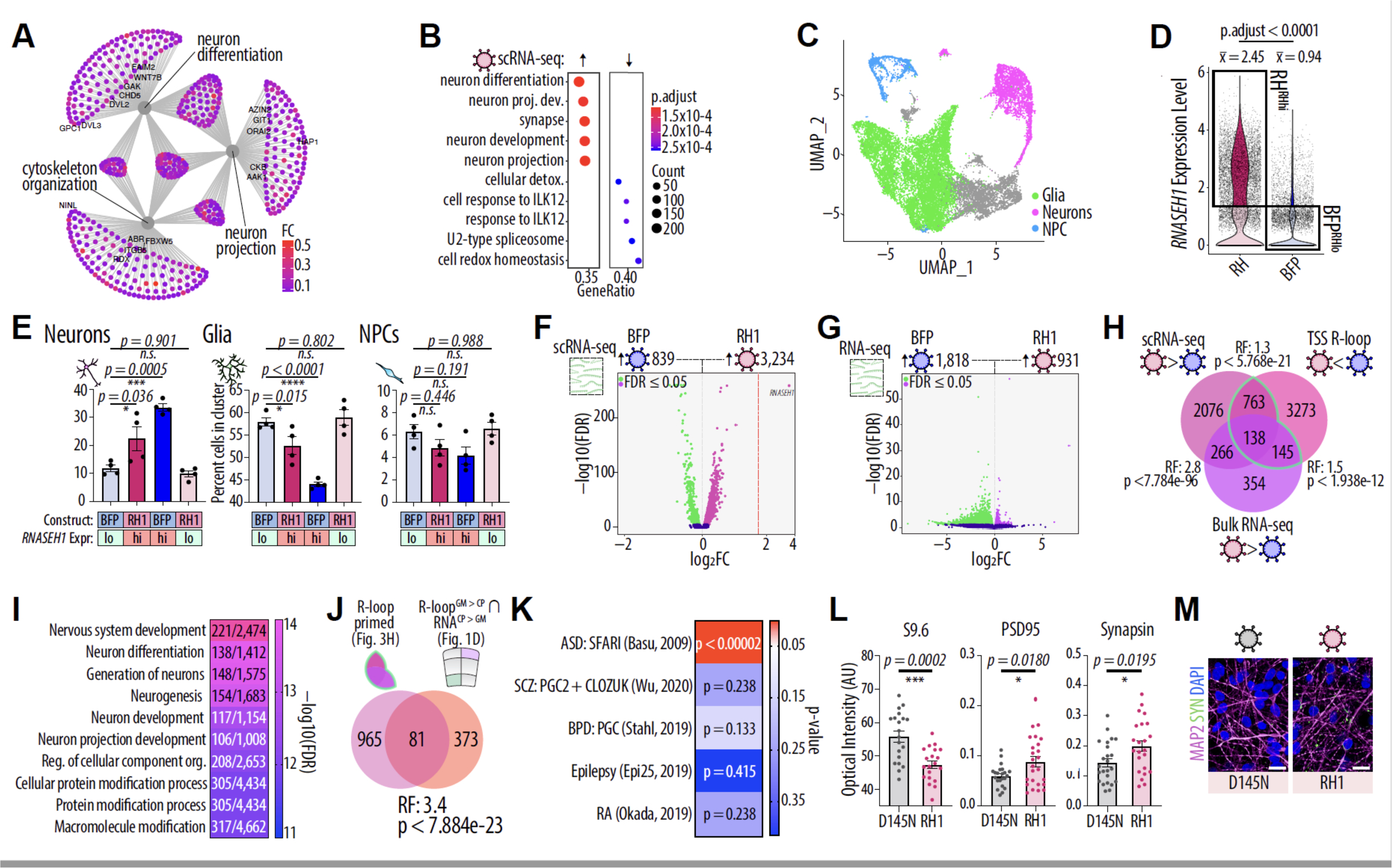
**Cell specifications and transcriptomic alterations in RH1 transgenic cultures.** **A, B)** Gene set enrichment analysis of scRNA-seq data from RH1ΔMLS and BFP control hiPSC-differentiated cultures. (A) Cnet plot and (B) dot plot showing significant gene ontologies up- (left) and down-regulated (right) following RH1ΔMLS transgene induction. Dot color indicates fold change (cnet plot) or adjusted p- value (dot plot), dot size indicates number of genes in category. n = 2 replicates from 2 cell lines/group. **C)** Combination of clusters into broad cell type classes, including glia (green), neurons (purple), and NPCs (blue). Gray clusters did not show canonical markers of these major neural cell types. **D)** RH1ΔMLS cells have significantly (p < 0.0001, Wilcoxin Rank Sum test) higher *RNASEH1* expression than BFP controls (mean normalized expression: RH1ΔMLS = 2.45, BFP = 0.94). Cells from each condition were split into two groups according to *RNASEH1* normalized expression value (> 1.25 = hi; ≤ 1.25 = lo) for downstream analyses. **E)** Bar charts showing difference in cell proportions across BFP or RH1ΔMLS transduced cells, stratified by *RNASEH1* expression level. There is a significant reduction in the proportion of glia (p = 0.015) and an increased proportion of neurons (p = 0.036) in RH1ΔMLS-*RNASEH1*^high^ relative to BFP-*RNASEH1*^low^ groups (Two-way ANOVA with Dunnett’s Multiple Comparison test; n = 2 replicates from 2 cell donors per group; error bars denote S.E.M.; *n.s.* = not significant.) **F, G)** Volcano plots showing differentially expressed genes in scRNA-seq (**F**) and bulk RNA-seq (**G**) from RH1ΔMLS hiPSC-differentiated cultures and BFP controls. Green dots indicate genes more highly expressed in control cells (FDR < 0.05, log_2_ fold change < 0) and purple/magenta dots indicate genes more highly expressed in RH1ΔMLS cells (FDR < 0.05, log_2_ fold change > 0). **H)** Venn diagram showing overlap of upregulated genes in RH1ΔMLS hiPSC-differentiated cultures from scRNA and bulk RNA-seq, and promoter-bound R-loops that are knocked down following RNase H1-transgene expression. RF = Representation factor, which signifies more overlap of the two gene sets than expected based on a background list of genes if > 1. P-values calculated via hypergeometric test. **I)** Gene ontology of genes with upregulated expression following promoter-bound R-loop loss in RH1ΔMLS hiPSC-differentiated cultures. Numbers within gene ontology images indicate number of genes in the enriched set that occur within the ontology class, legend denotes -log_10_ enrichment FDR. **J)** Venn diagram showing overlap of genes that have upregulated gene expression following promoter-bound R-loop loss in RH1ΔMLS hiPSC-differentiated cultures and R-loop “primed” genes in germinal matrix (Figure 1D). RF = Representation factor, p-values calculated via hypergeometric test. **K)** Heatmap indicating overlap of genes that have upregulated gene expression following promoter-bound R- loop loss in RH1ΔMLS hiPSC-differentiated cultures and disease risk genes. p-values calculated via hypergeometric test. **L)** Optical intensity from immunofluorescence images of the R-loop-specific antibody S9.6, and the synaptic proteins PDS-95 and synapsin, in RH1ΔMLS and RH1ΔMLS^D145N^ hiPSC-differentiated cultures. P-values calculated via two-tailed student’s t test; Error bars denote S.E.M.; n = 25 images per cell line / group. **M)** Representative immunofluorescence images of the synaptic protein synapsin (green) in RH1ΔMLS hiPSC- differentiated cultures and RH1ΔMLS^D145N^ controls, along with the nuclear marker DAPI (blue) and the neuronal marker MAP2 (magenta). Scale bar: 20 mm.

We identified 14 distinct cell clusters in our integrated scRNA-seq dataset using a graph-based clustering approach followed by Louvain modularity optimization (Supplementary Figure 5A). We then identified biomarkers within each cluster and defined its corresponding cell type using the Cell Type-Specific Expression Analysis (CSEA) tool [50] (Bonferroni adjusted p-value < 0.05, log_2_ fold change > 0; Supplementary Table 14). These annotations were confirmed using established cell type-specific marker transcripts (Supplementary Figure 5B) and clusters were grouped into the major divisions NPC, neuron, and glia, after removing cells not falling into these major categories (Figure 3C). The cell type heterogeneities and proportions observed in our culture system via scRNA-seq are consistent with previous studies employing this directed differentiation approach [51], which reflect the dynamic differentiation trajectory of hiPSC-NPCs into multiple neuronal subtypes together with non-neuronal cell types during neural development. We found no apparent changes to the number or proportions of cells in various stages of the cell cycle (BFP vs. RH1ΔMLS cells in phase, G1: p > 0.9, G2/M: p > 0.9, S: p > 0.9; Two-way ANOVA with Sidak’s Multiple Comparison test; Supplementary Figure 5C-D; Supplementary Table 15). Furthermore, the average expression of *RNASEH1* was not significantly different between glia, neurons, and NPCs in RH1ΔMLS cells (Two-way ANOVA with Tukey’s Multiple Comparison Test; glia vs. neuron: p = 0.3004; glia vs. NPC: p = 0.7761; neuron vs. NPC: p = 0.1260).

Interestingly, in BFP-transduced control cultures, the level of *RNASEH1* was significantly higher in neurons than in glia (p = 0.0001) and NPCs (p < 0.0001). This finding indicates a higher endogenous level of *RNASEH1* in neurons as compared to the other cell types (Supplementary Figures 5E-F), suggesting that R-loop regulation via RNase H1 is important for maintaining the neuronal lineage. To explore this further, we binned individual cells from our scRNA-seq assays based on the log_2_-normalized expression level of *RNASEH1* (high = *RNASEH1* > 1.25; low = *RNASEH1* ≤ 1.25) (Figure 3D, Supplementary Figure 5G-H). Interestingly, irrespective of the construct transduced, cells with high levels of *RNASEH1* expression were significantly more likely to differentiate into neurons (RH1ΔMLS: p = 0.036, BFP: p = 0.0005) than glia (RH1ΔMLS: p = 0.015, BFP: p < 0.0001; Two-way ANOVA with Dunnett’s Multiple Comparisons test) compared to BFP controls with low *RNASEH1* expression (Figure 3E). In contrast, RH1ΔMLS cells with low *RNASEH1* levels showed no significant difference from BFP controls with low *RNASEH1* expression, as it pertains to the proportion of cells that differentiated into glia (p = 0.802) or neurons (p = 0.901) (Figure 3E; Supplementary Figure 5I). Taken together, these findings strongly suggest that levels of *RNASEH1* expression in neural progenitor cells are associated with a differentiation shift towards the neuronal lineage.

Differential gene expression analysis identified 839 downregulated and 3,234 upregulated genes in RH1ΔMLS cells with high *RNASEH1* expression vs. BFP controls with low *RNASEH1* expression (FDR < 0.05, log_2_ fold change > 0; Supplementary Table 14) (Figure 3F). This general trend toward increased gene expression following *RNASEH1* transgene overexpression was also found within specific cell clusters (Glia: 562 down- and 2,778 up-regulated genes, Supplementary Table 16; Neurons: 225 down- and 335 up-regulated genes, Supplementary Table 17; NPCs: 77 down- and 226 up-regulated genes, Supplementary Table 18; Supplementary Figures 5J-L). We further confirmed these findings using bulk RNA-seq from RH1ΔMLS hiPSC-differentiated cultures and BFP controls (Figure 3G, Supplementary Table 19). We examined a potential link between altered gene expression after RH1ΔMLS expression and knock-down of promoter-bound R-loops. Indeed, there was a total of 1,045 genes showing upregulated expression via scRNA- and/or bulk RNA-seq together with a corresponding knockdown of the gene’s promoter-bound R-loop (p < 1.939e-10, representation factor = 1.2, hypergeometric test) (Figure 3H; Supplemental Table 20). Top gene ontology terms for these 1,045 *in vitro* R-loop primed genes following RNase H1 overexpression included synaptic signaling, nervous system development, and neuron projection development (Figure 3H). Importantly, a significant proportion of these *in vitro* R-loop primed genes were part of the list of 454 genes that were putatively primed by R-loops during neuronal differentiation in prenatal human brain *in vivo* (Figure 1D, Supplemental Table 21) (overlap n = 81, p < 7.884e-23, RF = 3.4; hypergeometric test) (Figure 3J). We conclude that during the process of neural progenitor differentiation *in vivo* and in the cell culture model, regulated expression for a subset of differentiation-sensitive genes is dependent on promoter-bound R-loops.

Next, we wanted to examine the disease-relevance of the collection of 1,045 genes defined by upregulated expression together with loss of the corresponding promoter-bound R-loop in RH1ΔMLS hiPSC-differentiated cells. Notably, this collection of R-loop primed genes was significantly enriched for risk loci associated with autism spectrum disorder [52]. This list includes, for example, *AUTS2*, a nuclear protein highly expressed in post-mitotic neurons of the prenatal brain [53], multiple cell adhesion molecules including *NRXN1*, *CDH7*, and *CDH8* [54] and ion channels including *CACNA1B* [55] (Supplementary Table 22). In contrast, common variants linked to schizophrenia risk [56–58], bipolar disorder [59], seizure disorder [60], or autoimmune disease [61] showed no significant enrichment for R-loop primed genes (Figure 3K; hypergeometric test).

Finally, we wanted to explore whether the observed differentiation shift toward the neuronal lineage, and the de-repression of neuronal transcripts following R-loop knockdown in our RH1ΔMLS cultures, wound be reflected in corresponding alterations at the protein level. Indeed, there was a significant upregulation of the synaptic proteins, PSD-95 (p = 0.018) and Synapsin (p = 0.020; Two-tailed student’s t test) in catalytically active RNase H1-overexpressing hiPSC neurons. In contrast, S9.6 immunoreactivity was significantly decreased (p = 0.0002) (Figure 3L, 3M).

### Contraction of the genome-wide R-loop space during neuronal differentiation is associated with deficits in functional and structural connectivity

Having shown that loss of promoter-bound R-loops in hiPSC-differentiated neural cultures results in hyperexpression of many autism and neurodevelopmental risk genes, we then asked whether these mechanisms are potentially detrimental for orderly neuronal development. To explore, we assessed spontaneous population-wide neuronal activity in our hiPSC-differentiated cultures using an Axion multielectrode array (MEA) (Figure 4A-B), which is broadly important for timed neurodevelopmental processes including migration, differentiation, circuit formation, and synaptic development and plasticity [62]. Neurons expressing the RH1ΔMLS transgene, in comparison to BFP control neurons, showed a significant reduction in neuronal activity at weeks 8 in culture (Figure 4C-E) (Two-way ANOVA with Sidak’s multiple comparisons test weeks 2-8; n spikes: p < 0.0001, n bursts: p < 0.0001, weighted mean firing rate: p = 0.0001; similarly, week 8 data was analyzed separately with a two-tailed unpaired student’s t test: number of spikes: p = 0.0298, number of bursts: p = 0.0066, weighted mean firing rate: p = 0.0216; n = 2 independent donor lines, Figure 4F). These alterations were highly specific to neuronal electrophysiological maturation, as migration and motility assays [63] did not reveal significant differences between RH1ΔMLS and BFP neurons (Supplementary Figure 5M-N).

**Figure 4.**
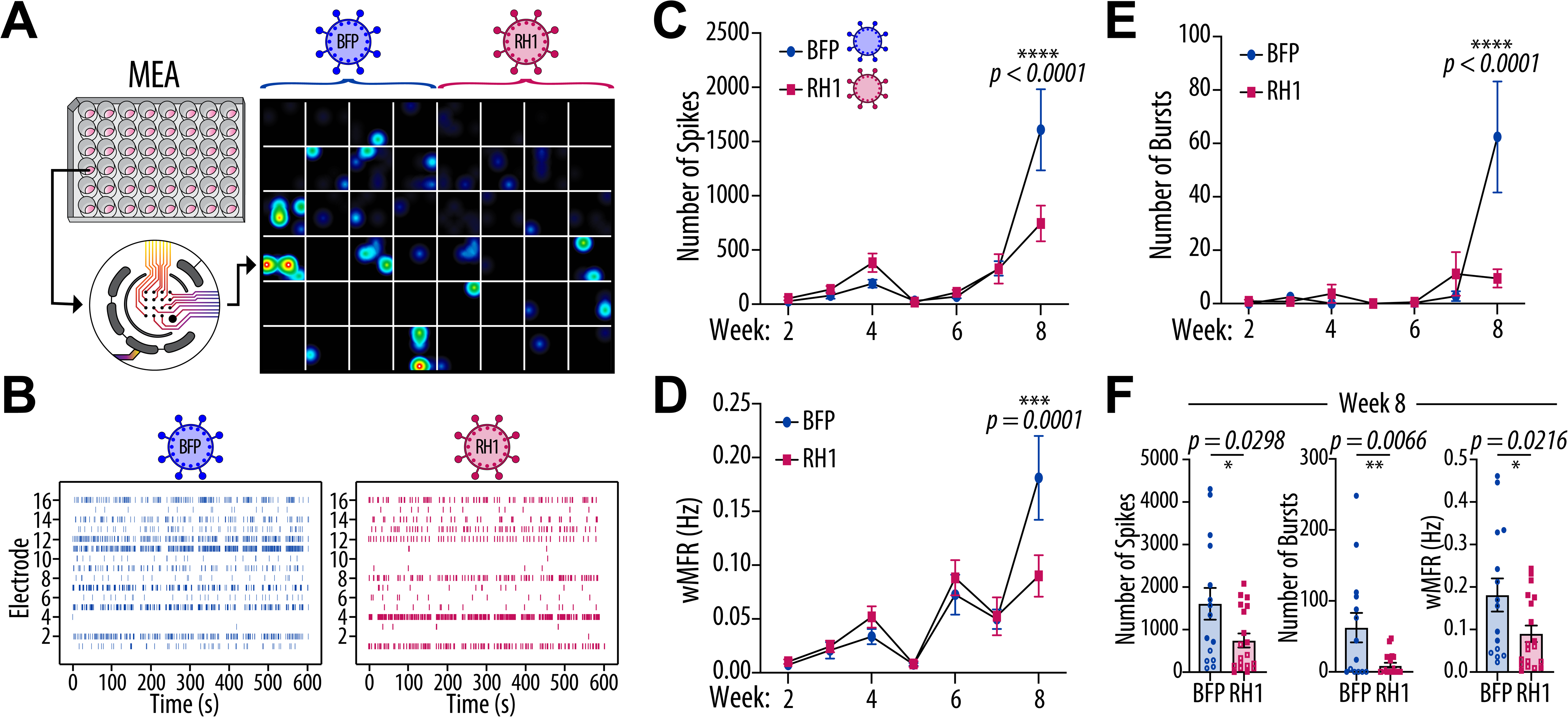
RNase H1-mediated R-loop loss during neuronal differentiation in vitro results in electrophysiological deficits. **A)** Top left: Electrophysiological recordings were taken with an Axion multielectrode array (MEA) system to assess population-wide neuronal activity. Schematic of the 48-well MEA plate used for electrophysiological recordings. Bottom Left: Schematic of an individual well within the MEA plate, containing 16 recording electrodes. Right: Screen shot of the MEA recording on week 8, with BFP-expressing hiPSC-neurons plated on the left half of the plate and RNase H1-overexpressing neurons on the right. **B)** Representative spike raster plots over the 10-minute recording at week 8 for BFP-expressing (left) and RNase H1-overexpressing (right) hiPSC-neurons. Each row denotes a single recording electrode within a single well of the 48-well MEA plate. **C-E)** The number of spontaneous spikes (**C**), weighted mean firing rate (wMFR) in Hertz (**D**), and number of bursts (**E**) were recorded weekly from BFP-controls and RNase H1 transgene expressing hiPSC-neurons from two donor lines. Data from the two donor lines were combined and a two-way ANOVA was performed with Sidak’s multiple comparisons test to test significant differences in each metric between weeks 2-8. Each condition contained at least 14-20 viable well replicates after removing wells that had less than 10 active recording electrodes. **F)** Statistics were additionally calculated for each electrophysiological metric at week 8 (unpaired two-tailed student’s t test) between RNase H1-transgene hiPSC-neurons and BFP controls. Individual donors represented with either filled in or hollow shapes.

Having shown that the RH1ΔMLS transgene is associated with defective functional connectivity *in vitro*, we then tested for evidence of defective structural connectivity *in vivo*. To this end, we used *in utero* electroporation (IUE) to overexpress RH1ΔMLS and GFP, or GFP and tdTomato as controls (Figure 5A), in the brains of embryonic day 15 (E15) C57BL/6 mice, targeting neural progenitor cells in the ventricular zone destined to differentiate into cortical pyramidal neurons [64, 65] (Figure 5B-4C). Expression of the RH1ΔMLS construct was confirmed via RT-qPCR of human *RNASEH1* in GFP+ neurons collected by fluorescence activated cell sorting (FACS) (p < 0.0001; Student’s Two-Tailed t test), while endogenous (mouse) *Rnaseh1* expression was unaffected (p = 0.9394) (Figure 5D). As expected for nuclear-targeted RH1ΔMLS, S9.6 immunoreactivity was diminished specifically in the nucleus of GFP+ neurons of mice electroporated with RH1ΔMLS and GFP (Figure 5E). Similarly, *RNASEH1* expression persisted in GFP+ cells in the medial prefrontal cortex (mPFC) of juvenile (P28) RH1ΔMLS mice, detected via fluorescence *in situ* hybridization (FISH) (Figure 5F). Quantitative 3D-image analysis with the Imaris Filament Tracer following an ultrafast optical clearing method (FOCM) [66] revealed that RNase H1 overexpression resulted in a significant reduction of dendrite complexity (Figure 5G, H) and dendritic spine density (Figure 5I, J; Apical spines: p < 0.01; basal spines: p < 0.05; Student’s Two-tailed t test) in pyramidal neurons of P28 mPFC, representing a developmental stage at which dendritic spine development robustly takes place in wildtype mouse cerebral cortex [67].

**Figure 5.**
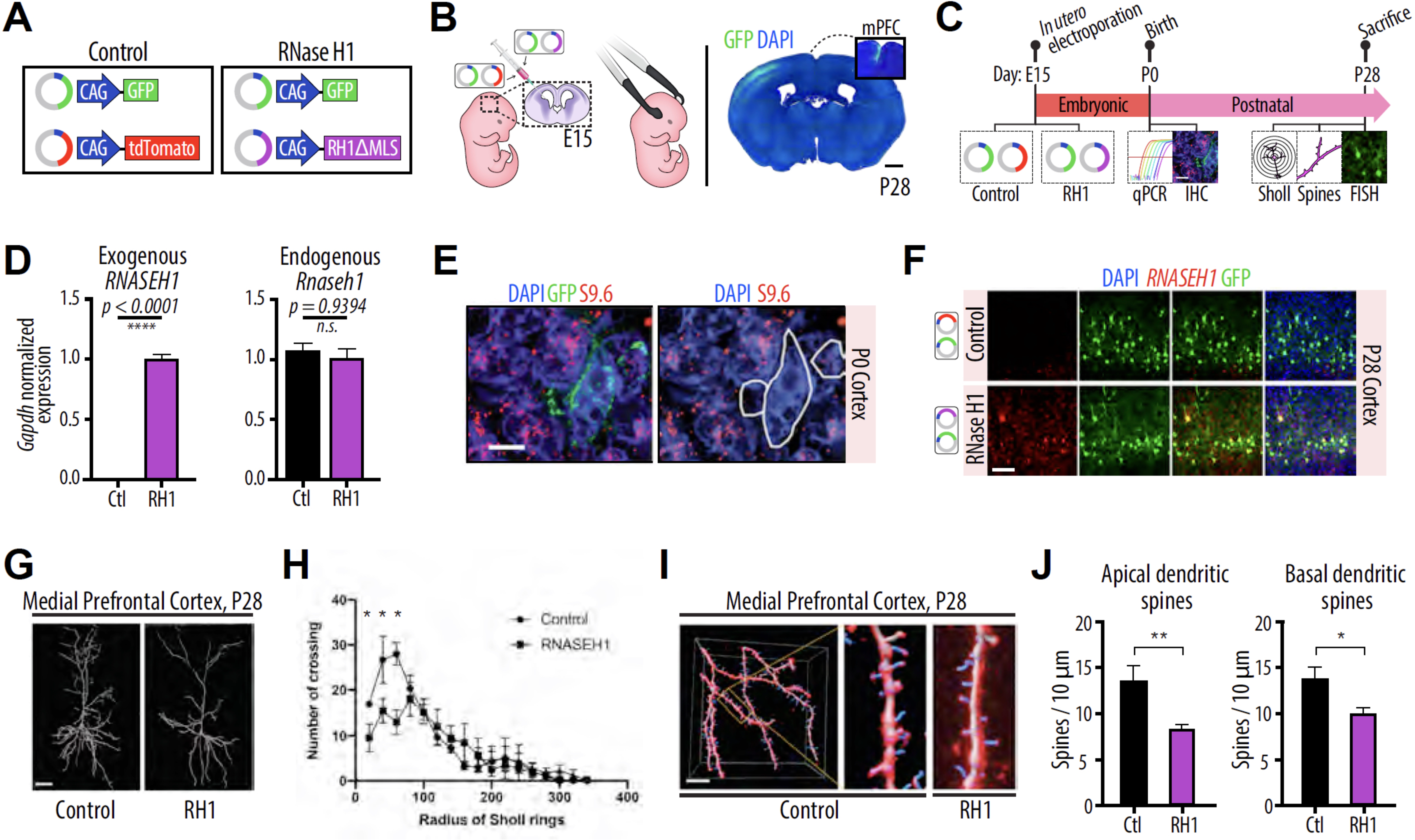
***RNase H1 overexpression leads to reduced dendritic complexity and spine density.*** Expression of the *RNASEH1*ΔMLS transgene, delivered via *in utero* electroporation into neural precursor cells of the ventricular zone in E15 mouse embryos, affects dendrite and spine morphology of cortical projection neurons. **A)** Expression plasmids of GFP combined with tdTomato are used as controls, while GFP combined with *RNASEH1*ΔMLS are used to destroy neural R-loops in mice during prenatal development. **B)** Overview of the *in utero* electroporation technique. TdTomato or RH1ΔMLS along with GFP reporters are transfected and electroporated unilaterally in E15 mouse ventricular zone. In adult mouse, P28, strong unilateral GFP expression is observed. Scale bar = 1 mm. **C)** *In utero* electroporation experimental timeline. **D)** Expression of exogenous (human) *RNASEH1* in RH1ΔMLS mice was confirmed in FAC-sorted GFP+ neurons, whereas no difference was observed for endogenous expression of (mouse) *Rnaseh1* between groups via qPCR (Normalized to *Gapdh* expression). Student’s two-tailed t test; n.s. = not significant. **E)** Expression of human RH1ΔMLS reduces S9.6 immunoreactivity in the nucleus of mouse neural cells at P0. **F)** FlSH images showing RNA expression of *RNASEH1* (red) in the P28 cortex of tdTomato controls and RH1ΔMLS mice, and colocalization of GFP (green) and *RNASEH1* in RH1ΔMLS cells. Scale bar = 100 mm. **G)** Representative images of (left) control and (right) RH1ΔMLS-expressing frontal pyramidal neuron dendritic trees at P28. **H)** Sholl analysis, showing reduced dendritic complexity in (left) control and (right) RH1ΔMLS-expressing pyramidal neurons at P28 (tdTomato control n = 3, RH1ΔMLS n = 4). **I)** Representative images of (left) control and (right) RH1ΔMLS frontal pyramidal neuron dendritic spines at P28. **J)** Apical and basal spine densities, assessed by Imaris tracing + FOCM (see text), are significantly decreased in RH1ΔMLS-expressing prefrontal pyramidal neurons. N = 3/group. *P < 0.05, **P < 0.01 determined by Two-Tailed Student’s t-test.

## DISCUSSION

Nascent pre-mRNAs, in addition to classical functions such as encoding proteins, also fulfill a chromatin regulatory role inside the cell nucleus [68]. This includes R-loops, defined as a DNA/RNA hybrid plus the displaced single-stranded sister DNA strand, which typically extend over 100-2,000 base pairs [69, 70]. R-loops are a specific molecular entity that is entirely separate from the much shorter, < 20 bp DNA/RNA hybrids generated in the ‘transcription bubble’ during the transcriptional process [71]. According to the present study, 62Mb, or up to 2% of the genome, is assembled in an R-loop configuration in the mid-gestational human prenatal brain,, which is of similar magnitude as comparable estimates for peripheral cells [5]. R-loop formation, topology, and kinetics are governed by multiple co-transcriptional mechanisms [72], and depend on the local chromatin environment to regulate activation, repression, or termination of gene transcription [73]. Further studies are required to elucidate the precise role of each of these mechanisms for local transcription in the developing brain.

Our genome-scale R-loop and transcriptome mappings in the prenatal brain, complemented by experimental, genome-wide reduction of the R-loop landscape in the NPC-to-neuron differentiation model *in vitro,* identified a subset of N = 456 developmentally regulated promoters that exist in the neural progenitor cell in an R-loop conformation. This subset of promoter-bound R-loops, while mostly enriched for neuron-specific and synaptic signaling genes, is at significant risk for premature expression when R-loop levels are experimentally decreased via overexpression of an RNase H1 transgene according to the findings presented hereoin. We therefore suggest that this group of promoter-bound R-loops underlies a poising mechanism for fine-tuning upregulated expression at later stages of neuronal development. Of note, the group of poised promoters carrying R-loops showed motif enrichment for repressor complexes previously linked to chromatin regulation involving R-loops, including SUZ12 Polycomb [42], RAD21 [74], and SIN3A histone deacetylase [75], that were sensitive to RNase H1-mediated R-loop removal. Therefore, the abnormal increase in gene expression following RNase H1-mediated R-loop removal in our NPC-to-neuron differentiation model *in vitro* could be associated with an imbalance of repressive and facilitative histone modification enzymes and chromatin remodeling complexes, a hypothesis that further supported the highly significant overlap (Figure 1E) between the group of R-loop bound poised promoters in human fetal brain described here, with a group of developmentally regulated neuronal genes in murine neural progenitors for which the poised status was defined by repressive and bivalent histone modifications in conjunction with RNA polymerase II pausing[25].

Importantly, the progressive deviation in electrophysiological activity in our hiPSC-derived neurons expressing the RNase H1 transgene resonates with findings that hiPSC-neurons derived from subjects diagnosed with autism spectrum disorder showed deficits in spikes and bursts after 50 but not 30 days of differentiation [76], and furthermore, reduced network connectivity has been reported in hiPSC-derived glutamatergic neurons carrying ASD-associated specific mutations [77], including the Fragile X Mental Retardation 2 gene (*AFF2*/*FMR2* [78]), and regulators of neuronal cell adhesion including *ASTN2* [79] and *NRXN1* [51]) which as shown here undergoes dynamic R-loop remodeling at the promoter together with transcriptional up-regulation during the course of neuronal differentiation (Table S22). Therefore, it is very likely that dysregulated expression and compromised poising status after R-loop ablation at the site of this group of genes contributes to the defective structural and functional connectivity in maturing neurons, as reported here.

## METHODS

### Cell culture

#### Generation and maintenance of hiPSC-derived NPCs

Human induced pluripotent stem cell (hiPSC)-neural progenitor cells (NPCs) were derived from hiPSCs as described previously [80]. Briefly, hiPSCs were generated via Sendai virus reprogramming and validated via karyotyping, gene expression, and immunocytochemistry as described elsewhere [81, 82]. hiPSC colonies were lifted with 1 mg/mL Collagenase in Dulbecco’s Modified Eagle Medium (DMEM) for 1-2 hours at 37°C and were transferred to a non-adherent plate, where they formed embryoid bodies (EBs) in suspension. EBs were cultured in suspension in a dualSMAD-inhibition media containing DMEM-F12-Glutamax (Invitrogen), 1X N-2 (Invitrogen), 1X B-27 (Invitrogen), 0.1 μM LDN193189 (Stegment), and 10 μM SB431542 (Tocris). After 7 days, EBs were plated on polyornithine/laminin-coated plates and cultured in N2/B-27 media containing 1 μg/mL laminin (Invitrogen). Neural rosettes were harvested by treating 14-day-old EBs with STEMdiff Neural Rosette Selection Reagent (Stem Cell Technologies) for 60 minutes at 37°C and were then plated onto polyornithine/laminin-coated plates and cultured with NPC medium containing DMEM/F-12, 1X N-2, 1X B-27-RA (Invitrogen), 1 μg/mL laminin and 20 ng/mL fibroblast growth factor 2 (FGF2).

hiPSC-derived NPCs were maintained at high density on Matrigel (Corning)-coated plates and were split 1:3 once a week with Accutase (Millipore) into NPC media containing DMEM/F-12 Glutamax, 1X N-2, 1X B-27, 20 ng/mL FGF2, and 1X Antibiotic/Antimycotic (Thermo Fisher Scientific). As reported previously, the doubling time of hiPSC-derived NPCs is 3.69 ± 0.05 days, with 62.3 ± 0.2% of cells in the G1 cell cycle phase, 24.6 ± 0.1% in S phase, and 6.5 ± 0.03% in the G2 phase [35].

#### Generation and maintenance of hiPSC-derived forebrain neurons

hiPSC-derived forebrain neurons were directly differentiated from hiPSC-NPCs as described previously [51]. Briefly, hiPSC-NPCs at passages between 9-11 were seeded at low density (e.g., 500k on a 6-well plate) on Matrigel-coated plates and were cultured for 6-8 weeks in neural differentiation medium, containing DMEM/F-12-Glutamax, 1X N-2, 1X B-27, 1X Antibiotic/Antimycotic, 20 ng/mL BDNF (Peprotech), 20 ng/mL GDNF (Peprotech), 1 mM dibutyryl-cyclic AMP (Sigma), 200 nM ascorbic acid (Sigma), and 1 μg/mL laminin. Half-media changes were performed every other day until cells were harvested. hiPSC-derived forebrain neurons reflect the dual-linage potential of NPCs and are therefore comprised of 70-80% BII- TUBILIN-positive neurons and 20-30% glial fibrillary acidic protein (GFAP)-positive glia. Of the 70-80% of neurons in the culture, approximately 30% express the GABAergic marker GAD67 while the rest show expression of the glutamatergic marker VGLUT1 [83].

### Postmortem Brain

Fresh and frozen de-identified prenatal autopsy samples with 4-72 hour postmortem intervals and lack of pathological abnormalities were collected by the laboratory of Dr. Nadejda Tsankova at the Icahn School of Medicine at Mount Sinai Pathology Department in accordance with the institutional policies and regulations, along with next-of-kin consent for the de-identified tissue to be used for research purposes. Exclusion criteria included maternal history of drug or alcohol abuse, autolysis interval > 72 hours, and RNA integrity values (RIN) of < 8.

Approximately 500 mg of tissue per sample was collected from second trimester prenatal human postmortem brain. Samples were dissected into two segments: 1) The germinal matrix (GM), situated next to the lateral ventricles, from which neurons and glia radially migrate toward the cortical plate (n = 3), and 2) the cortical plate (CP), which gives rise to the adult cerebral cortex (n = 3). As R-loops are very stable nucleic acid structures, more stable than dsDNA [84], they are not expected to degrade during tissue freezing and thus fresh autopsy and fresh frozen samples were combined in downstream DRIP-seq analyses.

### Animals

Timed-pregnant female C57/BL6J mice were purchased from The Jackson Laboratory for *in utero* electroporation and were housed, one to a cage, in a humidity- and temperature-controlled setting on a 12-hour light/dark cycle with *ad libitum* access to food and water. All experiments were performed in the Johns Hopkins University Brain Science Institute’s Behavioral Core according to the University’s Animal Care and Use Committee’s guidelines.

### S9.6 generation and validation

#### Hybridoma cell lines

The S9.6 antibody used for DRIP assays was generated in house from an S9.6-producing hybridoma cell line (ATCC; HB-8730). S9.6 cells were grown in serum-free media (ThermoFisher Scientific) and the antibody-containing supernatant was harvested by centrifuging cells at 1.5 X g for 5 minutes. The supernatant was purified via protein A affinity purification and size-exclusion, and was confirmed to be pure via SDS-PAGE, endotoxin-free (< 0.05 EU/mL), and protein identity was confirmed via mass spectrometry at the Albert Einstein College of Medicine Protein Production Facility. Purified S9.6 antibody concentration was determined via Bradford Assay, with a typical yield of ∼1 mg/mL.

#### Dot blot

The specificity of the S9.6 antibody for RNA/DNA hybrids was confirmed via dot blot using a protocol provided by Kerafast, which generates a commercially available S9.6 antibody. HPLC-purified oligonucleotides for the dot blot application were synthesized at IDT, and contained top and bottom strands of ssDNA and ssRNA (see Supplementary Table 23 for oligonucleotide sequences). Oligonucleotides (10 nM) were dissolved in annealing buffer (10 mM Tris, pH 8.0; 50 mM NaCl, 1 mM EDTA) to 100 nM and were heated at 95°C for 10 minutes, then cooled at room temperature for an additional 10 minutes. As a control, 15 units of RNase H1 (NEB, cat. no. #M0297) or 1 μg/μL RNase A (Life Technologies, cat. no. #AM2269) in low (0 mM) and high (500 mM) salt conditions were added to oligonucleotide pairs and incubated at 37°C for 1 hour to destroy DNA/RNA hybrids. All samples were purified via Phenol:Chloroform:Isoamyl alcohol (25:24:1, pH 8.0) (Invitrogen, cat. no. #15593-031) extraction and resuspended in 50 μL EB buffer (QIAGEN). A Hybond N+ positively charged nylon membrane (GE) was soaked in TBS-0.1% Tween 20 for 1 minute and allowed to dry for 10 minutes, then 2 μL of each oligonucleotide pair (DNA/RNA hybrid, dsRNA, and dsDNA) and negative controls were hybridized to the membrane by heating at 70°C for 2 hours. S9.6 antibody (1:1,000) was added and allowed to incubate overnight at 4°C, followed by a 1-hour anti-mouse secondary antibody incubation the following day (1:10,000). Antibody staining was visualized on a densitometer (BioRad). As shown in Supplementary Figure 1A, S9.6 binds specifically to the synthetic DNA/RNA hybrid exclusively, without binding to either dsDNA or dsRNA oligo pairs. Additionally, S9.6 signal was abolished after DNA/RNA hybrids were destroyed by RNase H1 or RNase A (0 mM NaCl) prior to membrane hybridization (Supplementary Figure 1A).

#### Western Blot

hiPSC-differentiated cultures were harvested with Accutase (Millipore) and pelleted with centrifugation for 5 mins at 1,000 G), then resuspended in 1X RIPA lysis buffer (Sigma) containing Complete Protease Inhibitor (Roche) and PhosSTOP phosphatase inhibitor (Sigma). Cells were then sonicated and centrifuged at 14,000 G for 30 min at 4°C. Total protein was separated on a 12.5% SDS-polyacrylamide gel (Bio-Rad) and transferred to a nitrocellulose membrane (Bio-Rad). The membrane was blocked with 4% bovine serum albumin (Sigma) in 1.0% Tween-20 (Sigma) in PBS. Following blocking, the membrane was incubated with primary antibody against RNase H1 (Invitrogen, cat. no. #PIPA530974, 1:1,000) and β-actin (Invitrogen, cat. no. #MA1-744, 1:5,000) overnight at 4°C with gentle agitation. The following day, the membrane was incubated with HRP-conjugated secondary antibodies (Sigma) for 2 hours at room temperature. The membrane was visualized with the Amersham ECL prime Western blotting kit (GE, cat. no. #RPN2232), using a Bio-Rad Gel Doc Imaging System.

High resolution (600 DPI) Western Blot images were converted to grayscale and quantified with Image J software [85]. The mean gray value from RNase H1 bands was divided by the mean gray value from β-actin bands, and fold change between treatment and control cells was calculated.

### RNase H1 transgene expression construct generation

#### Generation of RNASEH1ΔMLS plasmid

RNase H1 mRNA contains two in-frame AUGs, M1 and M27, which can either direct the translated protein to the mitochondria (beginning translation from M1) or to the nucleus (beginning translation from M27). As mitochondrial R-loops are essential for cell survival, we first used the QuikChange II Site-directed mutagenesis kit (Agilent, cat. no. #200523) according to the manufacturer’s instructions to remove the 78 bp mitochondrial localization signal (MLS) from RNase H1 cDNA (AddGene; pFRT-TODestGFP_RNAseH1; Plasmid #65784), using the primer sequences in Supplementary Table 24, RH1ΔMLS.

The plasmid was verified by Sanger sequencing at GENEWIZ and was used to generate both the TRE-*RNASEH1*ΔMLS-BFP lentivirus (for *in vitro* experiments) and the CAG- *RNASEH1* plasmid (used for *in vivo in utero* electroporation experiments) discussed below.

#### Generation of RNASEH1ΔMLS-D145N Plasmid

A single point mutation in the RNase H1 coding sequence renders the enzyme binding-competent but catalytically inactive. The aspartic acid (D) to asparagine (N) conversion at amino acid position 145 on the *RNASEH1*ΔMLS plasmid described above was achieved via site-directed mutagenesis using the primer sequences in Supplementary Table 24, *RNASEH1*Δ*MLS- D145N*, using Phusion DNA Polymerase (New England Biolabs, cat. no. #M0530L) and GC buffer under the PCR conditions found in Supplementary Table 24, *RNASEH1*Δ*MLS-D145N PCR Conditions*.

The plasmid was verified by Sanger sequencing at GENEWIZ and was used to generate both the TRE-*RNASEH1*ΔMLS-D145N-BFP lentivirus (for *in vitro* experiments) and the CAG-*RNASEH1*ΔMLS-D145N plasmid (used for *in vivo in utero* electroporation experiments) discussed below.

#### TRE-RNase H1ΔMLS-BFP lentivirus production

*RNASEH1*ΔMLS cDNA was subcloned into a doxycycline-inducible lentiviral backbone (under a TRE3G promoter), which also contained blue fluorescent protein (BFP) and puromycin resistance genes under an hPGK promoter. The plasmid was amplified and purified via Maxiprep (QIAGEN). Both TRE-*RNASEH1*ΔMLS-BFP and BFP-only control vectors were verified by Sanger sequencing at GENEWIZ.

Third-generation lentiviral packaging (VSVG) and envelope plasmids (REV, MDL), along with either TRE-*RNASEH1*ΔMLS-BFP, BFP-control, or reverse tetracycline-controlled transactivator (rtTA) plasmid, were polyethylenimine (PEI, Polysciences, cat. no. #23966-2)- transfected into HEK293T cells. Viral supernatant was collected 48 and 72 hours after transfection, centrifuged at 19,300 RPM for 2 hours at 4°C, resuspended in ice-cold DMEM (ThermoFisher), and stored in small aliquots at 80°C (with no aliquots freeze/thawing more than once). Viral titers were assessed with a qPCR Lentivirus Titration Kit (ABM) and functional titers were assessed by transducing iPSC-derived NPCs with serial dilutions of virus and quantifying BFP fluorescence intensity and RNase H1 protein levels by Western Blot.

#### CAG-HA-RNase H1ΔMLS plasmid production

The 5’ end of the *RNASEH1*ΔMLS cDNA was fused with an HA tag via restriction enzyme cloning, then subsequently subcloned into a CAG promoter-driven pCAGGS1 plasmid. As a negative control, green fluorescence protein (GFP) cDNA was restriction cloned into the CAG promoter-driven plasmid.

### RNase H1 transgene expression

#### Lentiviral Transduction in vitro

hiPSC-NPCs were transduced with either the TRE-*RNASEH1*ΔMLS-BFP, TRE- *RNASEH1*ΔMLS-D145N-BFP, or TRE-BFP viruses at a multiplicity of infection (MOI) of approximately 1.0. All constructs were co-transduced with the rtTA transactivator (Figure 2B). Virus was added to cells and allowed to incubate for 1 hour at 37°C, followed by spinfection at 1,000 G for 1 hour at room temperature with slow acceleration and deceleration. Cells were returned to 37°C and after 4 hours the media was changed (Day 0). Starting at Day 3, cells were fed every other day with NPC media containing 0.3 μg/μL puromycin. On Day 7, cells were split and replated at low density (e.g., 500,000 cells per well of a 6-well plate). The following day, cells were switched to neuron differentiation medium containing 0.05 μg/μL puromycin and 0.3 μg/μL doxycycline and were fed every 2-3 days for the course of their 6–8-week differentiation period.

#### In utero electroporation (IUE) in vivo

IUE targeting the cerebral cortex was performed using our previously published methods with minor modifications [86]. Pregnant C57/BL6 mice were anesthetized at embryonic day 15 (E15) by intraperitoneal administration of a mixed solution of Ketamine HCl (100 mg/kg), Xylazine HCl (7.5 mg/kg), and Buprenorphine HCl (0.05 mg/kg). After the uterine horn was exposed by laparotomy, the transgene-carrying plasmid (tdTomato or h*RNASEH1*) (1 µg/µl) together with CAG promoter-driven GFP expression plasmids (1 µg/µl) (molar ratio approximately 1:1) were injected into the lateral ventricles with a glass micropipette made from a micropillary tube (Narishige, cat. no. #GD-1), and electroporated into the ventricular zone of the medial prefrontal cortex (mPFC). For electroporation, electrode pulses (40 V; 50 ms) were charged four times at intervals of 950 ms with an electroporator (Nepagene, cat. no. #CUY21EDIT). All experiments were performed in accordance with the institutional guidelines for animal experiments from Johns Hopkins University.

#### Fluorescence-Activated Cell Sorting (FACS)

The mice subjected to IUE at E15 underwent whole brain extraction at P0. The cerebral cortex where GFP-positive neurons were localized was dissected using a stereomicroscope with a fluorescent flashlight (Night Sea, cat. no. #DFP-1). Cells were separated with a papain dissociation kit (Worthington, cat. no. #LK003150) with minor modifications [87]. For qPCR experiments, approximately 1.0-2.0 x 10^5^ GFP+ neurons per pup were collected by FACS into a 1.5 mL Protein LoBind Tube (Eppendorf, cat. no. #Z666505).

### DNA, Chromatin, and RNA extraction

#### DNA/RNA Immunoprecipitation (DRIP)

DNA/RNA Immunoprecipitation followed by Next Generation Sequencing (DRIP-seq) was performed as described in [4]. Genomic DNA was extracted from the specified tissue (∼500 mg) or cell type (roughly 5X10^6^ iPSC-NPCs or 1X10^6^ iPSC-neurons on a 6-well plate, enzymatically detached with Accutase (Millipore) and pelleted with centrifugation for 5 mins at 1,000 G), via incubation with 20% SDS and 20 mg/mL Proteinase K overnight at 37°C. After standard phenol/chloroform extraction, 3M NaOAc pH 5.2 and 2.4 volumes of 100% ice-cold EtOH was added and DNA was precipitated via gentle spooling with a pipette tip to avoid RNA contamination that can result from centrifugation. After three gentle washes with 70% EtOH and 1-2 hours of air drying, DNA was resuspended in TE and digested with a cocktail of restriction enzymes that avoid GC-rich regions (HindIII, EcoRI, XbaI, and SSPI) overnight at 37°C. DNA was again purified via phenol/chloroform extraction and ethanol precipitation, and complete digestion was confirmed on an agarose gel.

For immunoprecipitation of R-loops, a total of 13.2 μg of digested DNA (split into 3 tubes each containing 4.4 μg DNA) was combined with binding buffer (100 mM NaPO_4_ pH 7.0, 1.4 M NaCl, 0.5% Triton X-100) and 10 μL of 1 mg/mL S9.6 antibody was added to each tube. Samples were incubated with rotation overnight at 4°C. The next day, magnetic protein A/G beads (Thermo Fisher Scientific, cat. no. #88803) were washed with binding buffer 3 X 10 mins and 50 μL was added to each sample tube followed by a 2 hour incubation on a rotisserie shaker at 4°C. Bead:antibody complexes were washed 3 X 10 mins with binding buffer, then 250 μL of elution buffer (50 mM Tris pH 8.0, 10 mM EDTA, 0.5% SDS) and 7 μL of Proteinase K (20 mg/mL) was added to each tube and incubated at 55°C for 45 minutes with agitation to elute DNA from the S9.6 antibody. A standard phenol/chloroform extraction was performed, and DNA concentration and purity was assessed with a NanoDrop spectrophotometer. RNase A was added to samples (0.5 μL of 1 μg/μL stock) and incubated at 37°C for 1 hour. DNA was sheared with a Covaris S220 sonicator to a peak fragment size of 300 bp (duty cycle: 2%, incident power: 105 Watts, cycles per burst: 200, bath temperature: 4°C, processing time: 3 mins).

DRIP-seq was validated by pretreating hiPSC-derived NPC genomic DNA with RNase H1 and RNase A (in 0 mM NaCl) to destroy R-loops prior to sequencing. Consistent with previous reports, we observed an approximately 50% reduction in R-loop signal after RNase pretreatment [23] (Supplementary Figure 1D). We do not expect a 100% reduction in R-loops with RNase H1 and RNase A treatment alone, as some R-loops have been reported to be resistant to RNase degradation [88, 89]. This indicates that the DRIP-seq technique is able to map R-loops genome-wide, a large portion of which are sensitive to RNase pretreatment.

#### RNA Isolation and Purification

RNA was extracted from cells by adding 1 mL TRIzol (Invitrogen, cat. no. # 15596026) directly to the culture plate and scraping vigorously with a pipette tip. RNA was extracted with chloroform and precipitated with isopropyl alcohol, then treated with 2 U/10 μg RNA RNase-free DNase I (Invitrogen, cat. no. #AM222) at 37°C for 30 mins to remove DNA contamination. RNA was cleaned with acid phenol/chloroform and ethanol precipitated. Aliquots were stored at -80° prior to NGS library construction or RT-qPCR.

### Sequencing Data Generation and (RT)-qPCR

#### Next Generation Sequencing (NGS) Library Construction and Sequencing

NGS libraries from DRIP DNA and inputs were constructed according to the manufacturer’s instructions (KAPA Hyper Prep Kit: Roche, cat. no. #KK8502 kit and single-indexed adapters from Illumina, cat. no. #IP-202-1012 + IP-202-1024. Dual-indexed libraries were constructed with Roche, cat. no. #KK8722). Library fragment sizes were restricted to between 100-500 bp with a PippinHT instrument (Sage Science).

NGS libraries from total, purified RNA were constructed with the SMARTer Stranded RNA-seq Kit (Takara, cat. no. #634839) according to the manufacturer’s instructions for first-strand cDNA synthesis, purification of first-strand cDNA using SPRI beads, RNA-seq library amplification via PCR (with 12 PCR cycles for 10 ng of starting RNA), and SPRI cleanup.

All library sizes and concentrations used for molarity calculations were confirmed on an Agilent 2100 Bioanalyzer and a Qubit 4 fluorometer (ThermoFisher Scientific) with a Qubit dsDNA HS Assay kit (ThermoFisher Scientific, cat. no. #Q32854), respectively.

DRIP- and RNA-sequencing was performed on an Illumina HiSeq 2500 (single-indexed libraries). All sequencing was performed at the New York Genome Center, New York, NY, 2x150bp, to a minimum read depth of 50,000,000 reads. See Supplementary Table 2 for information about additional sequencing metrics.

### Bioinformatic analyses

#### DRIP-seq Bioinformatic Pipeline

DRIP-seq FASTQ files were aligned to GRCh37 with Bowtie2 [90]. Duplicates and reads with MAPQ score < 20 were removed with SAMtools [91], as well as the ENCODE blacklist [92] with BEDTools [93]. Following tag directory generation, peaks were called and annotated with HOMER [94] using an FDR threshold of 0.01 and style = histone against input samples. Standard quality control metrics were assessed with ChIPseeker [95], ChIPQC [96], NGS Plot [97], and deepTools [98].

Differential R-loop occupancy was calculated using the DiffBind package [99] normalized to the number of reads within consensus peaks (i.e., bFullLibrarySize = FALSE) when comparing prenatal CP & VZ/hiPSC-NPCs & hiPSC-NPC-differentiated cultures, and normalized to total aligned reads (i.e., bFullLibrarySize = TRUE) when comparing RNase H1- overexpressing cells and respective controls. When indicated, regions of interest were restricted to promoter regions (± 300 bp from TSS). DiffBind was used to generate volcano plots and heatmaps depicting differential R-loop occupancies. Differential peaks were annotated with ChIPseeker.

Gene ontology and ENCODE transcription factor binding site enrichment analyses were calculated with ShinyGO x0.61 [100] with an FDR threshold of 0.05 in the Biological Process category. Gene ontology for figures 1E and 1F were calculated from the top 1,000 (based on log_2_ fold change) genes within each set. For comparison of our DRIP-seq datasets with transcriptomic data, we analyzed RNA microarray data from 16-21 GW fetal brain obtained from the BrainSpan: Atlas of the Developing Human Brain resource [24] and calculated differentially expressed genes between the predominantly proliferative vs. post-mitotic tissues of the developing cerebral cortex. To anatomically align with our DRIP-seq data, the “CP” contained data from the BrainSpan inner and outer cortical plate regions, and the “GM” contained data from the BrainSpan inner and outer subventricular zone and the ventricular zone. We identified 1,562 significantly differentially expressed genes (DEGs) that were upregulated in the GM relative to the CP, and 1,263 DEGs upregulated in the CP relative to the GM (FDR ≤ 0.01; GM- upregulated DEGs = log fold change > 2; CP-upregulated DEGs = log2 fold change < -2). Similarly, we calculated differentially expressed genes between hiPSC-NPCs and hiPSC-neurons from our culture system [81], and found 1,074 genes upregulated in NPCs relative to neurons and 2,414 genes upregulated in neurons relative to NPCs (FDR ≤ 0.01; NPC-upregulated DEGs = log2 fold change > 1; Neuron-upregulated DEGs = log2 fold change < 1)).

Permutation analyses were performed with the RegioneR package [101] using a randomization function “resampleRegions,” with a background universe of all genes expressed in either fetal CP & GM or all genes expressed in hiPSC-NPCs & hiPSC-neurons, using 10,000 random permutations. Neuron-specific gene ontology analysis was calculated with SynGO [31].

R-loop feature distribution was calculated with ChIPseeker, using the default annotation criteria. To calculate promoter enrichment over GRCh37, promoter sequences (≤ 1kb from TSS) were downloaded from the UCSC Genome Browser and the overlap of the given gene set was tested against 10,000 random permutations (RegioneR) of all GRCh37 coordinates split into 1kb bins. DRIP-seq reads were visualized with the Integrated Genomics Browser program.

#### RNA-seq Bioinformatic Pipeline

After quality control with the FASTQC package and adapter trimming if necessary (Trimmomatic [102]), RNA-seq FASTQ files were aligned to GRCh37 using STAR [103] and uniquely mapped reads within genes were counted with FeatureCounts [104]. Ribosomal and mitochondrial RNA were removed prior to FeatureCounts with the BEDTools package. Genes with more than 10 counts in at least two samples were kept for downstream analysis. Sample-level quality control consisted of Principal Component Analysis and hierarchical clustering, both performed on regularized log transformed data. Differential gene expression was calculated with DESeq2 [45], α = 0.05. Data was visualized with R Studio, using the packages EnhancedVolcano and ComplexHeatmap.

#### scRNA-seq Bioinformatic Pipeline

Single cell RNA-sequencing was performed at the Mount Sinai Genomics Core. Samples were demultiplexed with the Seurat Package [105] and singlet cells with between 1,000 and 6,000 features and fewer than 25% reads mapping to the mitochondrial genome were kept for downstream analysis. Single cell transformation was applied with the SCTransform function in Seurat. Datasets from all samples were integrated using the FindIntegrationAnchors function in Seurat to identify anchors for merging the datasets with the IntegrateData command. The combined datasets were then scaled and two-dimensional t-distributed stochastic neighbor embedding (tSNE) was used to identify 14 clusters with the Seurat FindClusters function with the resolution parameter set to 0.35, determined via the Clustree package. RNA counts were log_2_-normalized with the NormalizeData function in Seurat and differential expression analysis was performed using these normalized counts. Positive cluster markers (Bonferroni adjusted p-value < 0.05, log_2_ fold change > 0) were identified and annotated with the CSEA tool [50] and confirmed with known cell type markers as described in the main text. Cells were divided into subgroups based on log_2_-normalized *RNASEH1* expression level with a threshold of 1.25, the level at which *RNASEH1* expression distribution between the control and treated groups overlaps. Differential gene expression was performed with the FindMarkers command of the Seurat package, using the “RNA” assay and “data” slot. Data was visualized with the EnhancedVolcano package. Gene Set Enrichment Analysis was performed and visualized with the ClusterProfiler package [106].

### Phenotypic assays

#### NPC migration assay analysis

To generate neurospheres, dissociated hiPSC-derived NPCs (n = 2 donors X 3 conditions: RH1ΔMLS, RH1ΔMLS^D145N^, and BFP) were grown in non-adherent plates in NPC media for 48 hours. Each well of a 96-well plate was coated in 2 μg/mL Matrigel two hours prior to manually picking individual neurospheres of roughly the same diameter and embedding into the Matrigel plates (n = 12 replicates per donor + condition). Neurospheres were centered within their wells and covered with an additional 25 μL of Matrigel. Brightfield images were taken on days 1 (1 day after embedding) and 3, and the total area covered by the neurosphere was calculated by tracing the perimeter of neurospheres using ImageJ software [85] on each day and measuring the area within. To calculate the distance traveled, the area (μm2) on Day 1 was subtracted from the area on Day 3, and an unpaired Student’s two-tailed t test was performed to detect significance between groups.

#### Immunostaining and analysis

For *in vitro* experiments, hiPSC-NPCs and neurons were grown on acid-etched coverslips in 48-well plates. For *in vivo* experiments, mice were perfused with 4% paraformaldehyde (PFA) as described above and coronal brain sections (40 μm) were collected using a freezing microtome and mounted onto Superfrost plus slides (Fisher). For staining with S9.6, brains were instead directly extracted and fixed in ice-cold 100% methanol and acetic acid for 2 hours. Cells *in vitro* were washed 1 X 5 min with ice-cold PBS then fixed with ice-cold 100% methanol (for nucleic acid, including R-loop, staining) or 4% PFA (for protein staining) for 10 minutes on ice, then washed 3 X 5 mins with ice-cold PBS. Cells or tissue sections fixed with PFA were further permeabilized for 15 or 30 minutes, respectively, at room temperature with 0.1% Triton X-100 in PBS (Thermo Fisher). Cells or tissue sections were blocked for 30 or 90 minutes, respectively, at room temperature in blocking solution (5% normal donkey serum with 0.01% Tween-20 in PBS), then incubated with primary antibody in blocking solution overnight at 4°C. Primary antibodies included RNase H1 (Invitrogen, cat. no. #PA530974, 1:250), Sox2 (Cell Signaling, cat. no. #2748, 1:500), MAP2AB (Sigma, cat. no. #M1406, 1:500), Nestin (Millipore, cat. no. #ABD69, 1:500), Synapsin (Millipore, cat. no. #574778, 1:500), PSD-95 (Invitrogen, cat. no. # 51-6900, 1:500), Fibrillarin (Abcam, cat. no. #ab4566, 1:500), and S9.6 (1:100, generated in house as described above). The following day, samples were washed 3 X 5 mins in ice-cold PBS and then incubated for two hours at room temperature with 1:500 secondary antibody in PBS, including Alexa donkey anti-rabbit 488, 568, 647 (Invitrogen) and Alexa donkey anti-mouse 488, 568, 647 (Invitrogen). In the last 10 minutes of secondary incubation, samples were counterstained with DAPI (Invitrogen) to visualize cell nuclei, then washed 3 X 5 mins in PBS at room temperature. Coverslips were mounted on Superfrost Plus slides (Fisher) and visualized on a ZEISS LSM780 confocal microscope.

When indicated (S9.6, Synapsin, PDS-95), optical intensity measurements were taken with ZEISS ZEN Black software, taking the highest optical density measurement within each 40 μm z-stack at 63X magnification. The average intensity value from five z-stacks within a single converslip was taken, and 20 total coverslips were used for the analysis. P-values were calculated with a two-tailed Student’s t test.

#### Dendrite Tracing/Sholl Analysis and Spine Density/Volume Analysis

Z-stack scanned images of GFP+ neurons in the mPFC of IUE mice were collected with a confocal microscope (ZEISS, LSM 700). All images were taken with 1024 x 1024 pixel resolution at a scan speed of seven per section. For Sholl analysis, images were taken with a 20X objective lens with a 0.50 µm interval for each image. Dendrite tracing was performed using Bitplane’s Imaris 9.3.1 software, specifically the Filament Tool’s Autopath algorithm. The method and specifications were adopted from a previous paper [107] with minor modifications. After manually selecting a region of interest, dendrites were specified as having a diameter between 0.6 and 11 μm. Seed points were then determined by the algorithm and set to the appropriate threshold to capture only the neuron present. Additionally, a region of 20 μm was designated around the soma to prevent extraneous tracing in the large area. After the algorithm’s execution, manual confirmation and any necessary edits were completed. To analyze the complexity of each neuron, Sholl analysis was performed via the Filament Sholl Analysis XTension for Imaris with Sholl sphere radius set to 20 μm with a maximum of 30 spheres.

For spine analyses, images focused on secondary apical or basal dendrites were taken by a 100X objective lens (with a 1.5X zoom factor) with a 0.24 μm interval for each image. Acquisition parameters were kept constant for all scans. Dendritic spine tracing was performed using the same Imaris Autopath algorithm and manual confirmation as dendrite tracing. The method and specifications were adopted from a previous paper [107] with minor modifications. Spines were designated with a minimum diameter of 0.4 μm and maximum length of 2.5 μm, the seed point threshold was set, and spines were digitally constructed. Per neuron, spine density was determined by averaging the number of spines along three different 10-15 μm dendrite segments. Mean spine volume was calculated by manually selecting the spines within the measured stretches and taking the automated calculation.

#### Multielectrode Array

hiPSC-derived NPCs from two control donor lines transduced with either BFP or RNase H1 virus were plated on a 48-well multielectrode array plate at a density of 80k cells/well (12 wells per donor/virus for a total of 48 wells). Each well of the MEA plate was coated with 2 μgLJmL−1 Matrigel 24 hours before seeding cells. One day following cell seeding, NPC media containing 1:1000 puromycin was replaced with neuron differentiation media containing 1:1000 Doxycycline and 1:5000 puromycin. Half media changes were performed every other day, and at least 24 hours before recording.

The Axion Maestro Multielectrode array system and AXIS software (Axion Biosystems) were used to perform 10-minute extracellular recordings of spontaneous network activity once a week for 8 weeks. Recordings were performed at 37°C and plates were first equilibrated for 10 minutes prior to recording. Data were sampled at a frequency of 12.5 kHz with a 200– 3,000 Hz single-order Butterworth band-pass filter and spike detection threshold 5.5 times the rolling standard deviation of the filtered field potential on each electrode. Wells with lifted cells by 8 weeks of culture were removed prior to analysis (7), as well as cells with fewer than 10 active electrodes (5). All conditions contained 7-10 viable well replicates.

Data from the two donor lines were combined and a two-way ANOVA was performed with Sidak’s multiple comparisons test in GraphPad Prism to test significant differences in the number of spikes, number of bursts, and weighted mean firing rates at weeks 2-8 in BFP- vs. RNase H1-transduced neurons. Similarly, an unpaired two-tailed t test was performed on week 8 data to detect significant differences in BFP- vs. RNase H1-transduced neurons. Spike train raster plots were generated with the Matplotlib function eventplot on MEA spike files from a single representative well from each condition.

#### Fluorescence In Situ Hybridization

Coronal mouse brain slices (20 μm) were collected using a freezing microtome and mounted onto Superfrost plus slides (Fisher). The slides were processed per the RNAscope Multiplex Fluorescent v2 protocol (Advanced Cell Diagnostics). Briefly, the tissue sections were treated with hydrogen peroxide (protease digestion was not performed) and incubated with probes targeting *Rnaseh1* (ACD, cat. no. #532251) mRNA for 2 hours at 40°C. Amplifier sequences were polymerized to the probes and treated with Opal 650 dye. The slides were cover slipped with DAPI Fluoromount (Southern Biotech). Imaging was performed on a ZEISS LSM780 confocal microscope.

## ACKNOWLEDGEMENTS

We thank all members of the Akbarian and Brennand laboratories for constructive comments and discussions. Bibi Kassim, Natalie Barretto, Samuel Powell, Esther Cheng, Yuto Hasegawa, and Sasha Layne for laboratory assistance. The New York Genome Center, Kristin Beaumont and the Mount Sinai Genomics Core Facility, and Prashant Singh at the Roswell Park Genomics Shared Resource for sequencing support. John Greally, Julie Nadel, Achla Gupta, and the Albert Einstein College of Medicine Proteomics Core for assistance with S9.6 antibody generation and purification. Zhiping Weng, Kaili Fan, Kiran Girdhar, and Gabriel Hoffman for productive discussions. We also thank Frédéric Chédin and Lionel Sanz for generously sharing reagents and the DRIP-seq protocol.

## DATA AND CODE AVAILABILITY

Data will be made available on Synapse.org.

## AUTHOR CONTRIBUTIONS

Experiments conducted by E.A.L. (hiPSC differentiations, *RNASEH1* cloning and lentiviral construction/transduction, DRIP-seq, RNA-seq, MEA, ICC, qPCR, Migration assay, WB, dot blot), A.S. and A.F. (IUE, spine density and Sholl analyses, *in vivo* IHC/qPCR), B.J. (FISH), A.H. (qPCR, optical intensity and image analyses), M.B.F. (hiPSC-neuron differentiation and ICC), K.J.B. (hiPSC/NPC generation), and N.M.T. (prenatal brain dissection). Bioinformatic analysis conducted by E.A.L. (DRIP-seq, RNA-seq, scRNA-seq), A.P.J. (scRNA-seq), C.D., S.E.G., J.E.E., M.E., and W.L. Research supervised by S.A., K.J.B., N.M.T., A.K., B.T., and L.S. Study and experiments conceived by E.A.L., S.A., and K.J.B. Figures designed by E.A.L. and S.A. Paper written by E.A.L., S.A., and K.J.B., with contributions from all co-authors.

## SUPPLEMENTARY FIGURES

Supplementary Figure 1. Validation of the S9.6 Antibody and DRIP-seq technique.

A) Dot Blot: Synthetic oligonucleotides (dsDNA, DNA/RNA, dsRNA) are hybridized to a positively charged membrane and S9.6 is allowed to bind (top). S9.6 only binds to DNA/RNA hybrids, which is abolished by pretreatment of RNase H1 or RNase A in low salt conditions (bottom).

B) Staining with the S9.6 antibody (red), the nucleolar marker fibrillarin (green), and DAPI (blue) in iPSC-derived NPCs and neurons shows expected staining profile, which is primarily located within nucleoli and surrounding the cell nucleus in mitochondrial-rich regions. Scale bar (top to bottom right) 50 m, 20 m, 50 m.

C) DRIP-seq Workflow: Genomic DNA is fragmented with a cocktail of restriction enzymes, incubated with S9.6 overnight, and R-loop/S9.6 complexes are captured with magnetic beads and unbound DNA is washed away. S9.6-bound DNA is eluted and made into high-quality next generation sequencing libraries for deep sequencing.

D) DRIP-seq following RNase treatment or with RNA alone: hiPSC-derived NPC DNA is treated with RNase H1 and RNase A prior to DRIP-seq and shows an expected ∼50% reduction of R-loops (left two bars). NPC RNA is extracted and taken through the DRIP-seq protocol instead of genomic DNA, and results in a negligible number of peaks called (rightmost bar). MiSeq platform, n = 1 sample per validation condition.

Supplementary Figure 2. DRIP-seq bioinformatic workflow and QC.

**A)** *In vivo* and *in vitro* samples from the present study and GSE70189 were aligned to GRCh37 via Bowtie 2 [90], converted to BAM files with duplicate reads and reads with MAPQ score < 20 removed via SAMtools [91], the ENCODE blacklist [92] was removed via BEDtools [93], and peaks were called and annotated using HOMER software with an FDR threshold of 0.01 [94] with input normalization. Downstream quality control and processing of DRIP-seq peaks utilized the software packages ChIPseeker [95], deepTools [98], and DiffBind [99].

**B-E)** We identified (mean ± SEM) 52,686 ± 11,840 DRIP-seq peaks in GM (peak detection FDR < 0.01) and 70,199 ± 8,043 peaks in CP. R-loop peaks have a median (μ_1/2_), range, and average (X = mean ± SEM) size of: μ_1/2_ = 630 bp, [373 bp – 24,003 bp], and X = 1,062 ± 85 bp in prenatal GM. μ_1/2_ = 641 bp, [368 bp – 27,462 bp], and X = 969 ± 32 bp in prenatal CP. Bp = basepairs, kbp = kilobasepairs. R-loops covered 1.84% ± 0.51% of the GM genome (spanning 57.6 ± 15.9 Mbp) and 2.18% ± 0.30% (spanning 68.5 ± 9.54 Mbp) of the CP genome (GM n = 3; CP n = 3). No statistically significant differences were found in prenatal GM or CP in number of DRIP-seq reads (B), number of R-loop peaks (C), size of R-loop peak (D), or percent of genome occupied by R-loops (E). See main text for details; Significance calculated with a one-way ANOVA with Tukey’s Multiple Comparison test. *n.s.* = not significant; error bars denote SEM.

Supplementary Figure 3. Individual R-loop dataset overlaps with prenatal brain RNA-seq.

**A)** Top: The number of overlapping genes between each individual GM-enriched R-loop dataset and CP- upregulated genes (BrainSpan RNA microarray data) (red bar) was tested for significance by performing 10,000 random permutations of the overlap of GM-enriched R-loops with a “background universe” of any expressed gene in the prenatal cortex (gray bars comprising normal distribution), blue bar = significance threshold (α = 0.05). Remaining graphs, top to bottom, similarly summarize permutation results for each individual CP-enriched R-loop dataset ∩ CP-upregulated genes, each individual GM-enriched R-loop dataset ∩ GM-upregulated genes, and each individual CP-enriched R-loop dataset ∩ GM-upregulated genes. Positive Z-scores indicate overlap of the target gene sets are higher than expected by chance.

**B)** Permutation testing as described above for (top to bottom) NTera2-enriched R-loops ∩ CP-upregulated genes, K562-enriched R-loops ∩ CP-upregulated genes, NTera2-enriched R-loops ∩ GM-upregulated genes, and K562-enriched R-loops ∩ GM-upregulated genes. * = data from GSE70189 [3].

Supplementary Figure 4. RNase H1 transgene and in vitro expression.

**A)** TRE3G-*RNASEH1*ΔMLS-PGK-BFP-PuroR lentiviral construct.

**B)** *RNASEH1* expression is significantly higher in RH1ΔMLS and RH1ΔMLS^D145N^ hiPSC-differentiated cells and NPCs. **** = p < 0.0001.

**C-E)** DRIP-seq confirms R-loop reduction in RH1ΔMLS cells relative to controls over genes (C) and promoter-regions (D) genome-wide, while R-loop levels are not altered in mitochondrial genes in RH1ΔMLS cells.

**F)** R-loop peaks have a median (μ_1/2_), range, and average (X = mean ± SEM) size of: μ_1/2_ = 694 bp, [316 bp – 58,071 bp], and X = 1,856 ± 36.79 bp in BFP-expressing hiPSC-differentiated cells (n = 3). μ_1/2_ = 631 bp, [401 bp – 68,044 bp], and X = 1,337 ± 170 bp in RH1ΔMLS^D145N^ differentiated cells (n = 2). μ_1/2_ = 626 bp, [402 bp – 53,255 bp], and X = 1,387 ± 41.16 bp in RH1ΔMLS differentiated cells (n = 3). Kbp = kilobasepairs.

**G)** No statistically significant difference was found in the number of DRIP-seq reads between BFP and RH1ΔMLS differentiated cells. RH1ΔMLS^D145N^ differentiated cells had significantly more DRIP-seq reads than BFP (p = 0.0271) and RH1ΔMLS (p = 0.0090).

**H)** RH1ΔMLS cells contained significantly less, (mean ± SEM) 21,528 ± 940, R-loop peaks compared to BFP (32,893 ± 1,035, p = 0.0061) and RH1ΔMLS^D145N^ (51,129 ± 3,153, p = 0.0001) differentiated cells.

RH1ΔMLS^D145N^ neurons had significantly more R-loop peaks relative to BFP-expressing (p = 0.0012) differentiated cells.

**I)** The genome span occupied by R-loops was significantly reduced in RH1ΔMLS differentiated cells, with (mean ± SEM) 29.9 ± 1.8 Mbp (0.95% ± 0.06%) of the genome covered, relative to BFP (61.1 ± 3.1 Mbp, 1.95% ± 0.10% of genome, p = 0.0012) and RH1ΔMLS^D145N^ (67.8 ± 4.5 Mbp, 2.16% ± 1.53% of genome, p = 0.0008) controls. No significant difference was found between BFP and RH1ΔMLS^D145N^ controls. See Supplementary Table 2 for complete description of DRIP-seq data; Significance calculated with a one-way ANOVA with Tukey’s Multiple Comparison test*. n.s.* = not significant; error bars denote SEM.

**J-K)** Volcano plots displaying the differential genome-wide R-loop peaks (DRIP-seq) between hiPSC-differentiated cells expressing RH1ΔMLS (n = 3) vs. BFP (n = 3) (J) or RH1ΔMLS^D145N^ (n = 2) (K) (FDR < 0.05, pink dots) after 6 weeks in culture.

Supplementary Figure 5. scRNA-seq in RNase H1 transgene-expressing and control cells.

**A)** Clustering of integrated scRNA-seq data reveals 14 distinct cell clusters.

**B)** Violin plots of expression of NPC (*PCNA*, *MKI67*, *CCNB1*), neuronal (*RBFOX3*, *MAPT*, *SYP, DCX*, *CRABP1*, *SYN1*), and glial (*AQP1*,*AQP4*, *ITGA6*) marker gene expression in each cell cluster, used to define broad cell types. Feature axis on log scale.

**C)** Bar chart displaying percent of cells in G1, G2/Mitosis, and S cell cycle phases within BFP and RH1ΔMLS groups. n = 2 replicates each from 2 donors/group; 2-way ANOVA with Sidak’s Multiple Comparison test; n.s. = not significant.

**D)** Distribution of cells in each scRNA-seq cluster in G1, G2/M, and S phases of the cell cycle in both BFP and RH1ΔMLS groups.

**E)** Normalized *RNASEH1* expression within all clusters in BFP-expressing and RH1ΔMLS groups.

**F)** Average normalized *RNASEH1* expression in glial, neuronal, and NPC clusters in RH1ΔMLS (left) and BFP (right) groups. Two-way ANOVA with Tukey’s Multiple Comparisons test; n = 2 replicates each from 2 donors/group; n.s. = not significant.

**G)** Violin plot displaying the normalized *RNASEH1* expression value within each cell; BFP and RH1ΔMLS groups were further subdivided based on their normalized *RNASEH1* expression level (high = *RNASEH1* > 1.25; low = *RNASEH1* ≤ 1.25). Total number of cells within each subgroup indicated at top.

**H)** Normalized *RNASEH1* expression within all clusters in RH1ΔMLS-*RNASEH1*^high^ (top left), BFP- *RNASEH1*^high^ (bottom left), BFP-*RNASEH1*^low^ (top right), and RH1ΔMLS-*RNASEH1*^low^ (bottom right) subgroups.

**I)** Percent of cells in each unannotated cluster in BFP-*RNASEH1*^low^, RH1ΔMLS-*RNASEH1*^high^, BFP- *RNASEH1*^high^, and RH1ΔMLS-*RNASEH1*^low^ subgroups. Unless otherwise indicated, differences in cell proportions relative to controls (BFP-*RNASEH1*^low^) were not significant (Two-Way ANOVA with Dunnett’s Multiple Comparisons test).

**J, K, L)** Volcano plots showing differential gene expression (scRNA-seq) between RH1ΔMLS-*RNASEH1*^high^ and BFP-*RNASEH1*^low^ cells, refined to glia (**H**), neurons (**I**), and NPCs (**J**) (Bonferroni adjusted p-value < 0.05). Green dots indicate genes more highly expressed in BFP-*RNASEH1*^low^ cells (log2 fold change < 0) and purple/magenta dots indicate genes more highly expressed in RH1ΔMLS-*RNASEH1*^high^ cells (log2 fold change > 0).

**N)** The area gained in RNase H1-, RNase H1-D145N, and BFP-overexpressing hiPSC-neurospheres (n = 2 independent donor lines). Dots indicate independent wells from each group (n = 25). Distance traveled = Area (μm^2^) Day 3 – Day 1: BFP = 2.28 X 105 μm^2^, SEM = 9.2 X 103 μm^2^; RNase H1 = 2.37 X 105 μm^2^, SEM = 12.1 X 103 μm^2^; unpaired student’s two-tailed t test, p = 0.5925. N.s. = not significant, error bars denote SEM.

Representative images of migrating neurospheres 3 days after cell culture plating, and depiction of area calculation strategy in ImageJ. Scale bar: 200 m.

