## Supplementary figures and images for "R-loop landscapes in the developing human brain are linked to neural differentiation and cell-type specific transcription"

### Supplementary Figure 1

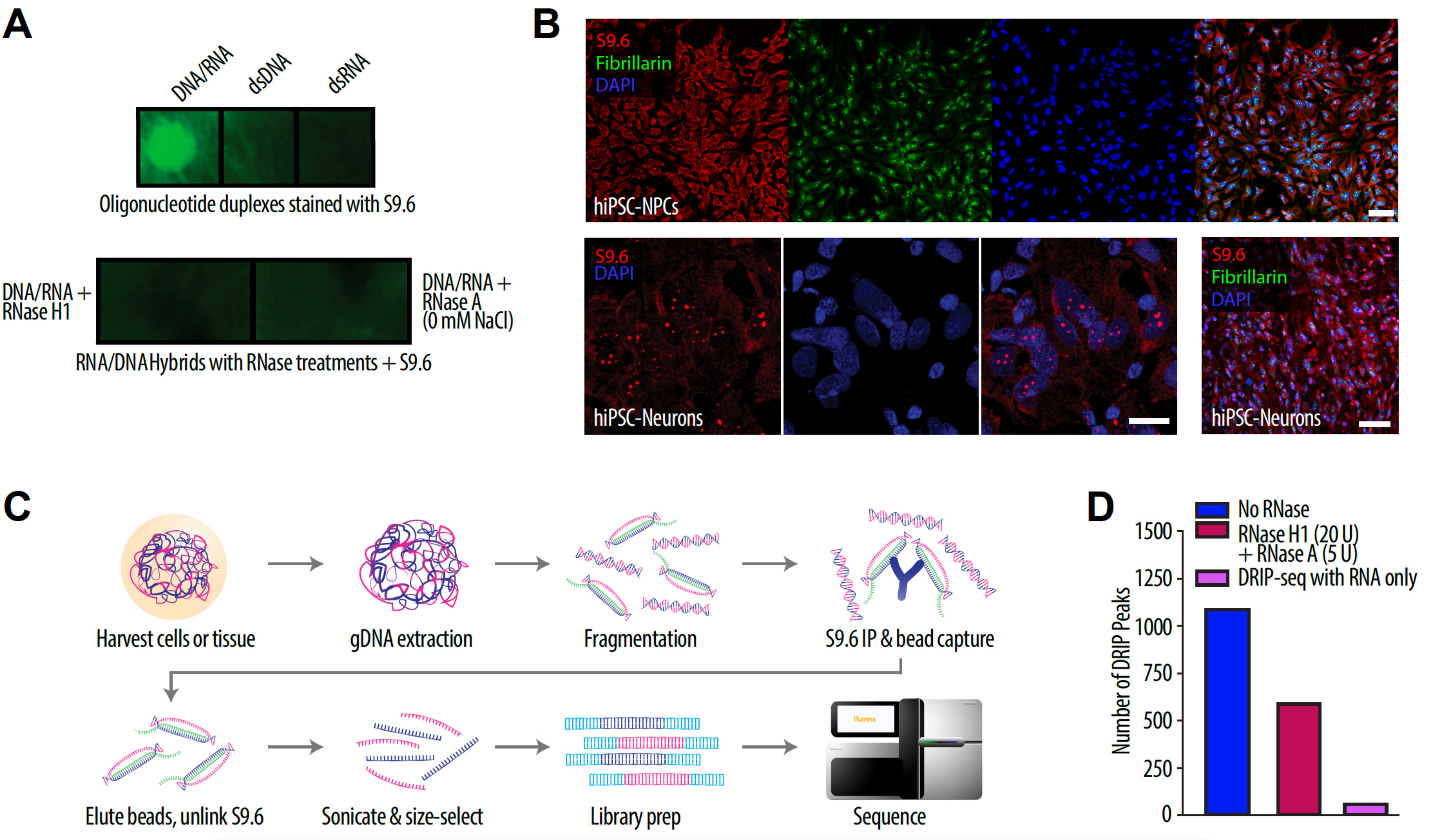

### Supplementary Figure 2

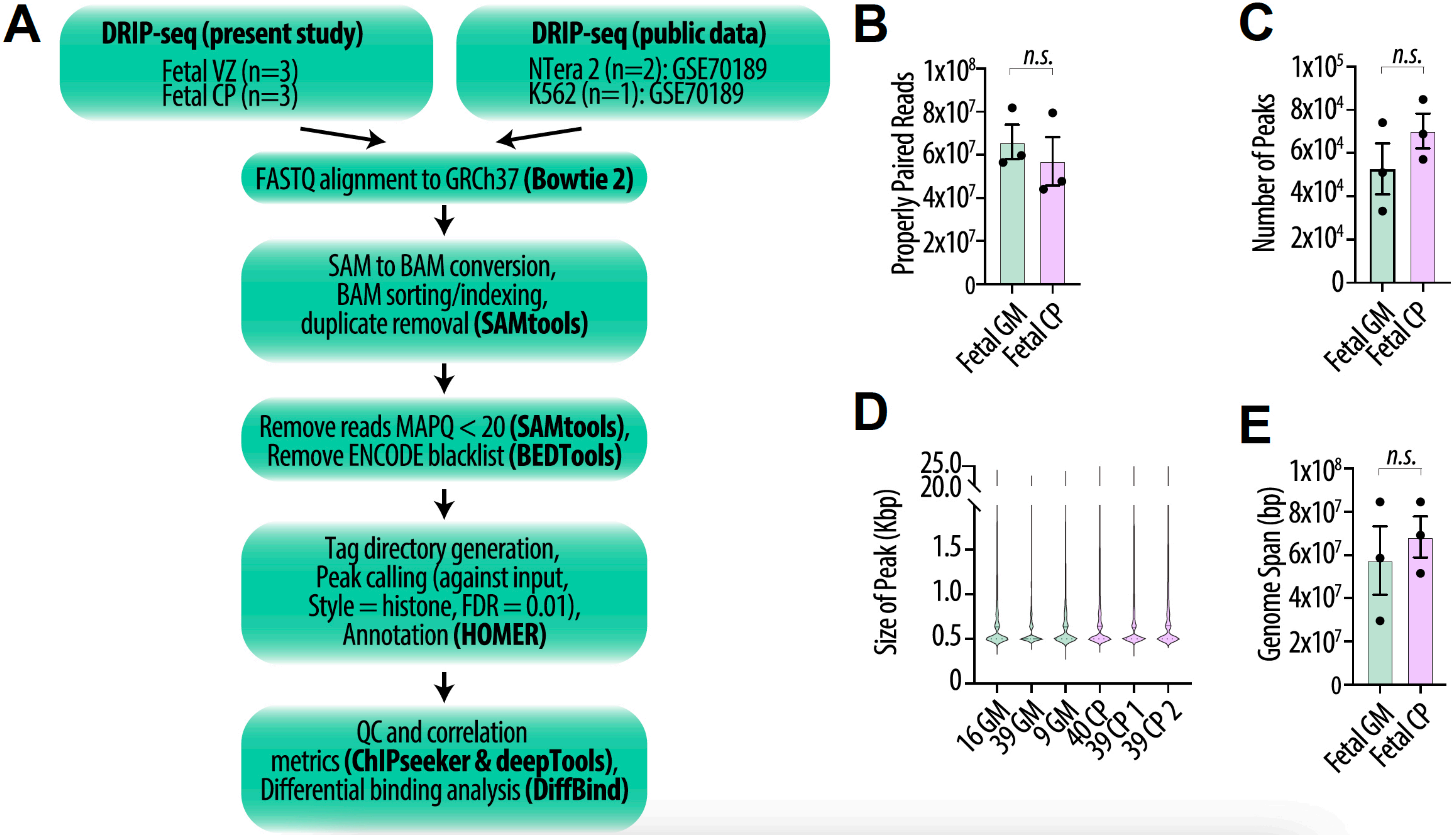

### Supplementary Figure 3

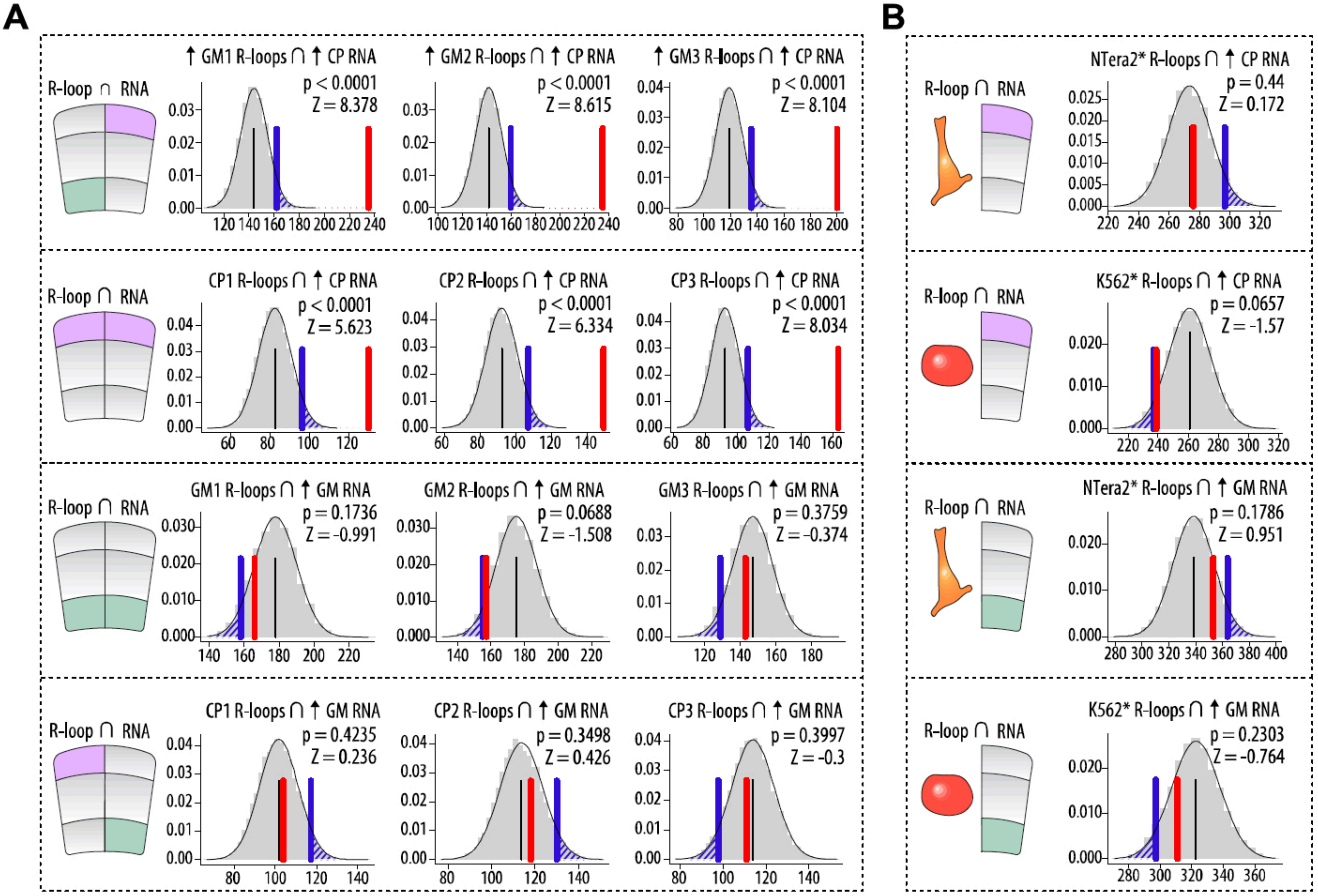

### Supplementary Figure 4

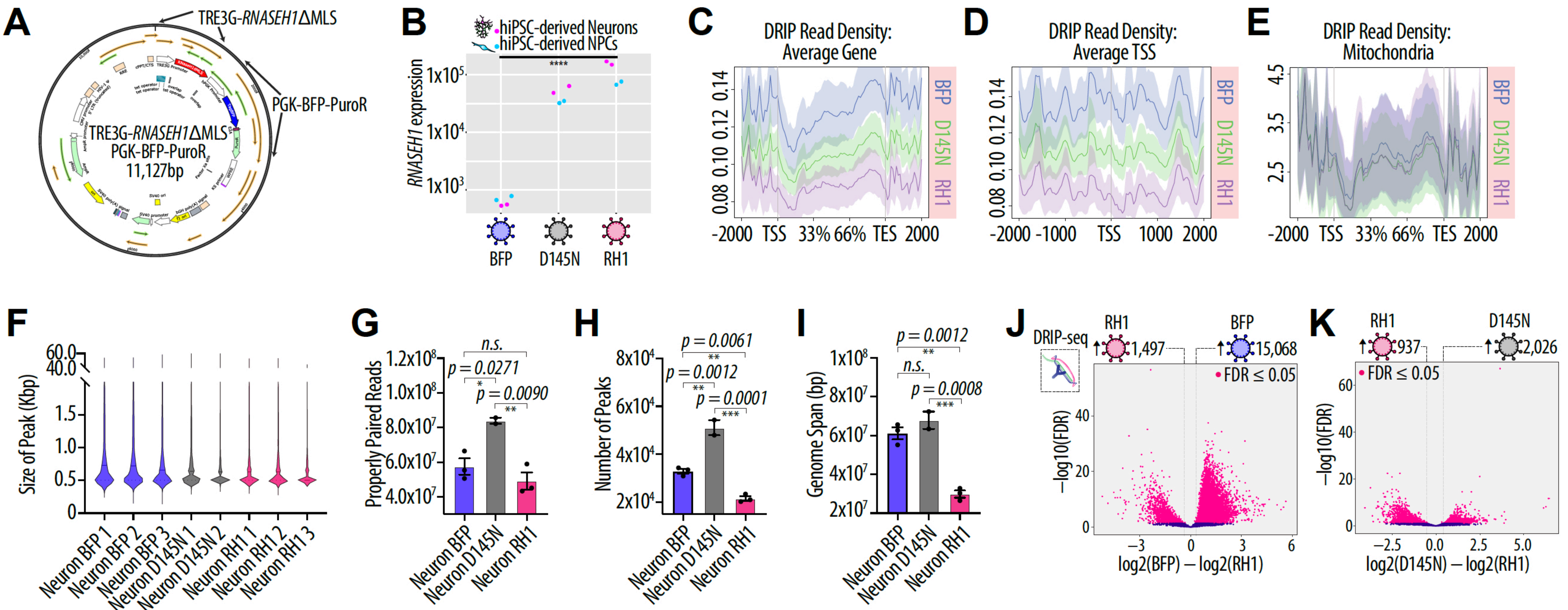

### Supplementary Figure 5

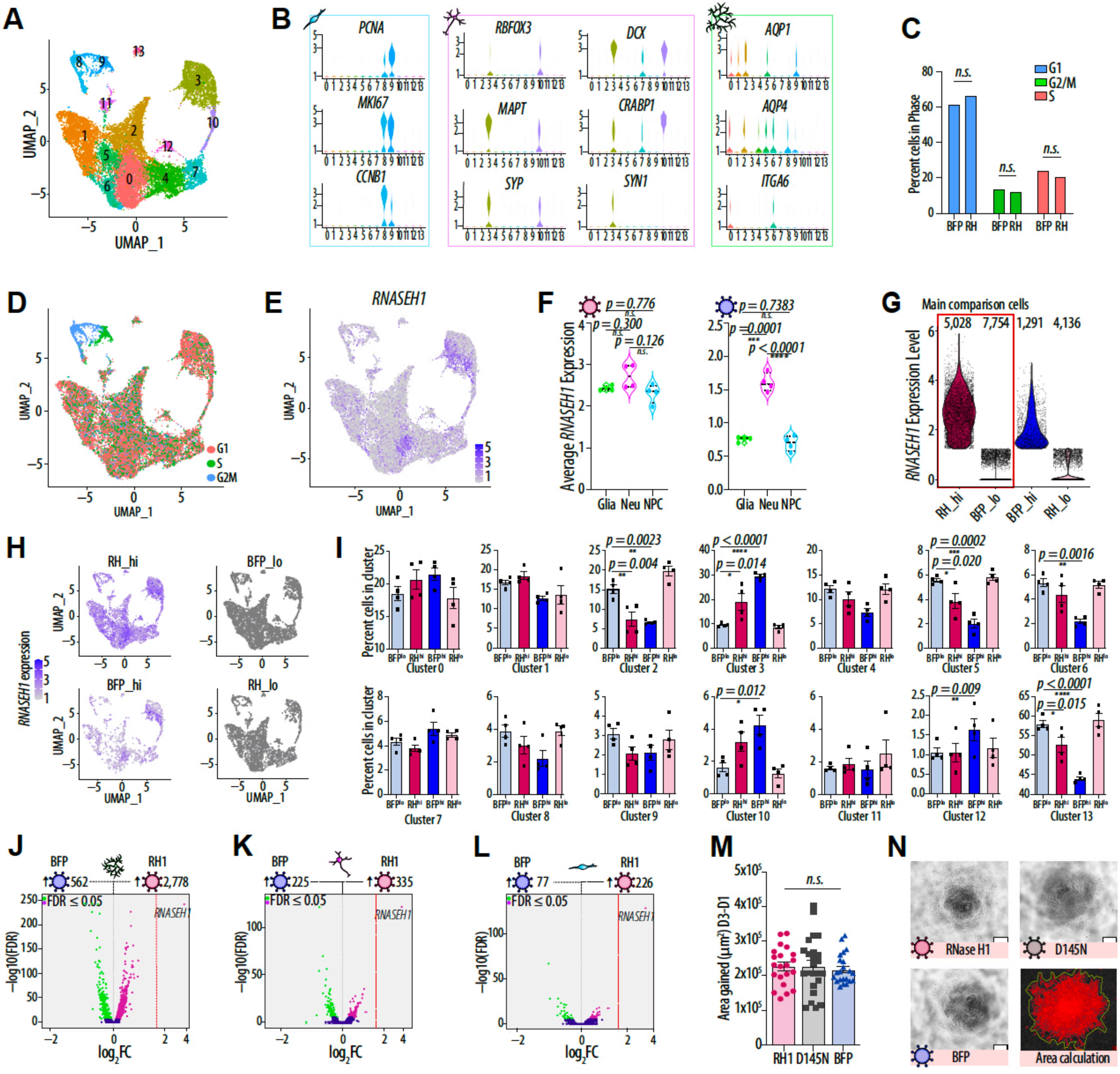
